# Targeting the Immune-Fibrosis Axis in Myocardial Infarction and Heart Failure

**DOI:** 10.1101/2022.10.17.512579

**Authors:** Junedh M. Amrute, Xin Luo, Vinay Penna, Andrea Bredemeyer, Tracy Yamawaki, Gyu Seong Heo, Sally Shi, Andrew Koenig, Steven Yang, Farid Kadyrov, Cameran Jones, Christoph Kuppe, Benjamin Kopecky, Sikander Hayat, Pan Ma, Guoshai Feng, Yuriko Terada, Angela Fu, Milena Furtado, Daniel Kreisel, Nathan O. Stitziel, Chi-Ming Li, Rafael Kramann, Yongjian Liu, Brandon Ason, Kory J. Lavine

**Affiliations:** Center for Cardiovascular Research, Division of Cardiology, Department of Medicine, Washington University School of Medicine, Saint Louis, MO, 63110, USA; Genome Analysis Unit, Amgen Discovery Research, Amgen Inc., 1120 Veterans Blvd, South San Francisco, CA, 94080, USA; Department of Cardiometabolic Disorders, Amgen Discovery Research, Amgen Inc., 1120 Veterans Blvd, South San Francisco, CA, 94080, USA; Mallinckrodt Institute of Radiology, Washington University School of Medicine, Saint Louis, MO, 63110, USA; Institute of Experimental Medicine and Systems Biology, RWTH Aachen University, Aachen, Germany; Department of Internal Medicine, Erasmus Medical Center, Rotterdam, The Netherlands; Division of Cardiothoracic Surgery, Department of Surgery, Washington University School of Medicine, Saint Louis, MO, 63110, USA; The Jackson Laboratory, Bar Harbor, ME 04609, USA; Department of Pathology and Immunology, Washington University School of Medicine, Saint Louis, MO, 63110, USA; Department of Genetics, Washington University School of Medicine, Saint Louis, MO, 63110, USA; McDonnell Genome Institute, Washington University School of Medicine, Saint Louis, MO, 63110, USA; Department of Developmental Biology, Washington University School of Medicine, Saint Louis, MO, 63110, USA

**Keywords:** myocardial infarction, fibroblast activator protein, macrophages, C-C chemokine receptor 2, interleukin 1 beta, fibrosis, Cellular Indexing of Transcriptomes and Epitomes by sequencing.

## Abstract

Cardiac fibrosis is causally linked to heart failure pathogenesis and adverse clinical outcomes. However, the precise fibroblast populations that drive fibrosis in the human heart and the mechanisms that govern their emergence remain incompletely defined. Here, we performed Cellular Indexing of Transcriptomes and Epitomes by sequencing (CITE-seq) in 22 explanted human hearts from healthy donors, acute myocardial infarction (MI), and chronic ischemic and non-ischemic cardiomyopathy patients. We identified a fibroblast trajectory marked by fibroblast activator protein (FAP) and periostin (POSTN) expression that was independent of myofibroblasts, peaked early after MI, remained elevated in chronic heart failure, and displayed a transcriptional signature consistent with fibrotic activity. We assessed the applicability of cardiac fibrosis models and demonstrated that mouse MI, angiotensin II/phenylephrine infusion, and pressure overload models were superior compared to cultured human heart and dermal fibroblasts in recapitulating cardiac fibroblast diversity including pathogenic cell states. Ligand-receptor analysis and spatial transcriptomics predicted interactions between macrophages, T cells, and fibroblasts within spatially defined niches. CCR2^+^ monocyte and macrophage states were the dominant source of ligands targeting fibroblasts. Inhibition of IL-1β signaling to cardiac fibroblasts was sufficient to suppress fibrosis, emergence, and maturation of FAP^+^POSTN^+^ fibroblasts. Herein, we identify a human fibroblast trajectory marked by FAP and POSTN expression that is associated with cardiac fibrosis and identify macrophage-fibroblast crosstalk mediated by IL-1β signaling as a key regulator of pathologic fibroblast differentiation and fibrosis.

## Introduction

Cardiac fibrosis is a major contributor to life-threatening arrhythmias, heart failure progression, and mortality. While present in nearly all forms of cardiac disease, cardiac fibrosis is prominent in patients who suffer a myocardial infarction (MI) and those with ischemic cardiomyopathy, where coronary artery stenosis and/or occlusion leads to loss of viable myocardium^12^. Despite advancements in clinical management, MI and ischemic cardiomyopathy are among the leading causes of heart failure, associated morbidity, and death worldwide^3, 4^ and continue to impose staggering financial burden on healthcare systems^3, 5–8^.

Strategies to therapeutically target cardiac fibrosis have remained elusive. Recent advances in CAR T-cell technology have provided a much needed breakthrough and indicated that selectively removing activated cardiac fibroblasts that express fibroblast activator protein (FAP) is sufficient to ameliorate fibrosis without impacting the structural integrity of the heart in pre-clinical mouse models^9, 10^. While these studies suggested that targeted fibroblast subset depletion may be therapeutically achievable and challenged the dogma that cardiac fibrosis is irreversible, many questions remain to be answered. The precise fibroblast populations that are responsible for fibrosis in the human heart and mechanisms that orchestrate their emergence and maturation are unknown. Furthermore, it is not clear which of the many available pre-clinical animal and *in vitro* experimental models ^11–16^ best recapitulate human cardiac fibrosis.

The advent of high-throughput sequencing technologies have enabled high resolution profiling of healthy and diseased tissues to discover novel cell types and behaviors^17, 18^. Numerous single-nucleus (snRNA-seq) and single-cell RNA sequencing (scRNA-seq) studies have profiled the healthy human and failing human heart^19–23^ and observed expansion of immune cells^21^ and fibroblast diversification in the failing heart^20, 23^. However, they did not identify human cardiac fibroblast populations that drive fibrosis. A drawback of the existing studies is the inclusion of only RNA expression information and heavy reliance on single nucleus RNA sequencing datasets, which restrain high-resolution cell phenotyping due to limited transcriptome coverage.

To identify fibroblast cell states, associated transcriptional signatures, cell surface protein markers, and signaling events that drive fibrosis in the human heart, we performed Cellular Indexing of Transcriptomes and Epitomes by sequencing (CITE-seq)^24^ in human hearts from non-diseased donors, chronic ischemic, and non-ischemic cardiomyopathy patients. We utilized a panel of 279 antibodies to phenotype cell types and states that emerge in disease to comprehensively define human cardiac fibroblasts diversity and to uncover cell surface protein targets for diagnostic and therapeutic applications. Moreover, we leveraged scRNAseq data from multiple mouse cardiac injury models and human fibroblast cell culture systems to assess the suitability of experimental systems to recapitulate human cardiac fibroblasts diversity in health and disease. We employed cell interaction analysis and spatial transcriptomics in human MI tissue and identified immune-fibroblast neighborhoods within areas of fibrosis that were predicted to be driven by IL-1β signaling from macrophages to activated fibroblasts. Finally, we demonstrated that IL-1 signaling to fibroblasts is essential for cardiac fibrosis in mice and elucidated that IL-1 signaling coordinates the emergence and maturation of a cardiac fibroblast trajectory marked by FAP and POSTN. These findings establish a causal role for inflammation in pathological fibroblast fate specification, maturation, and fibrosis.

## Results

### CITE-seq defines the cellular landscape of human myocardial infarction and heart failure

To characterize the transcriptional and surface proteomic landscape of human MI and heart failure, we performed CITE-seq in human left ventricle (LV) specimens obtained from 6 healthy donors, 4 acute MI patients (<3 months post-MI), 6 ischemic cardiomyopathy (ICM, >3 months post-MI) patients, and 6 non-ischemic cardiomyopathy (NICM, idiopathic dilated cardiomyopathy) patients (**Fig. 1A**). Donor hearts were obtained from brain dead individuals with no known cardiac disease and normal LV function. Acute MI, ICM, and NICM specimens were obtained at the time of left ventricular assist device (LVAD) implantation or heart transplantation. We included samples from males and females (**Table 1**). Hematoxylin and eosin (H&E) staining showed marked inflammation in acute MI specimens and robust fibrosis in ICM and NICM specimens (**Fig. 1B**).

**Figure 1.**
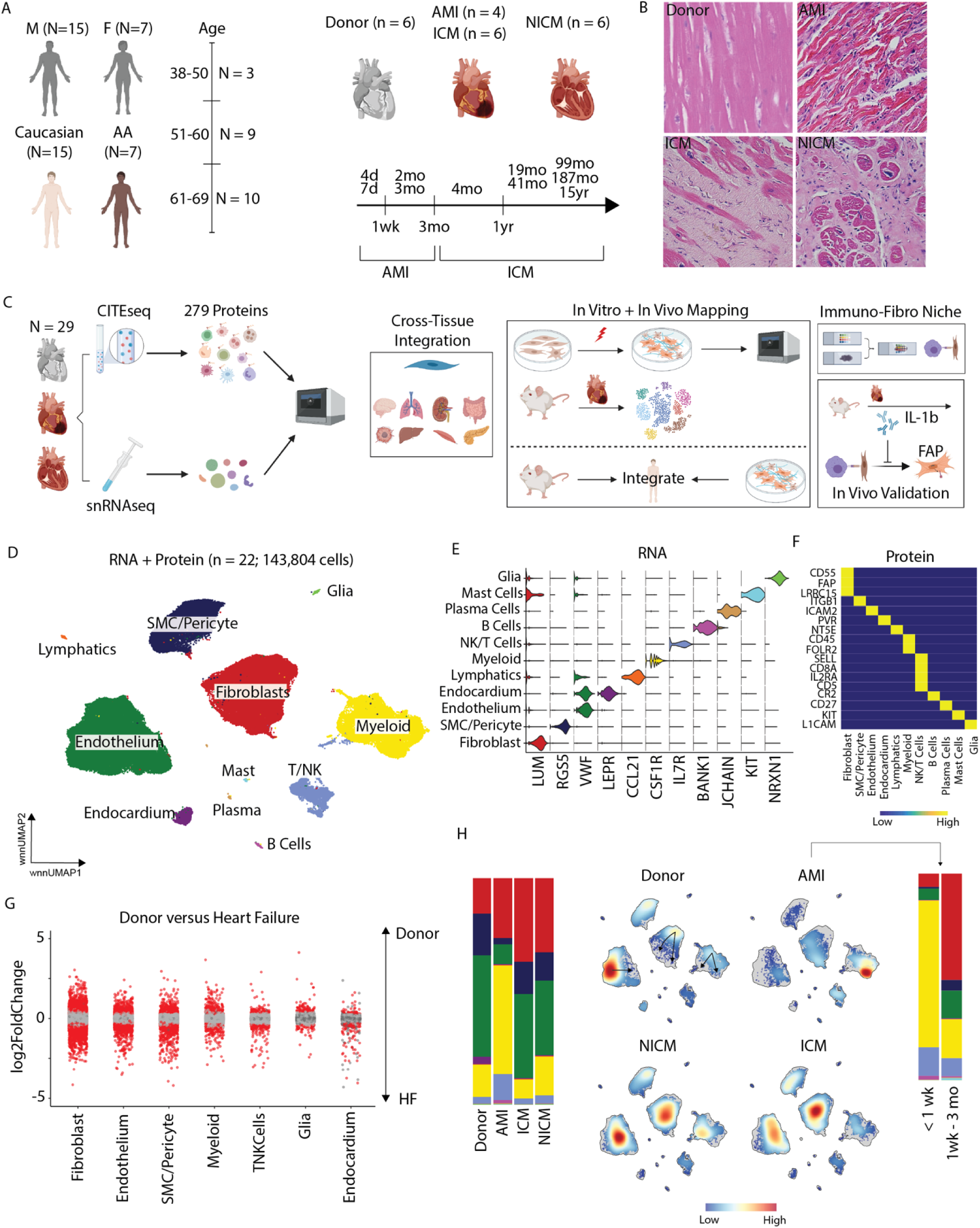
Study design and integrated clustering across donor and HF samples. (A) General patient demographics. (B) Representative H&E sections from donor, acute MI, ischemic cardiomyopathy, and non-ischemic cardiomyopathy. (C) Study design. (D) UMAP embedding plot with weighted nearest neighbor clustering of CITE-seq data from 22 patients and 143,804 cells. (E) Violin plot for canonical marker genes for cell types. (F) Heatmap of marker proteins for cell types. (G) Pseudobulk differentially expressed genes between donor and HF samples in major cell types. Red dots indicate statistically significant genes (log2FC > 0.58 and adjusted p-value < 0.05). (H) Cell cluster composition across conditions (left), cell density in four conditions (middle), and cell cluster composition in acute MI patients split by MI time.

CITE-seq using 279 surface antibodies was performed on transmural myocardial specimens obtained from the LV anterior wall (**Fig. 1C, Supplementary Fig. 1A**). After pre-processing and application of quality control filters, we obtained high-quality transcriptomic and proteomic reads in 143,804 cells (**Supplementary Fig. 1B-C**) and performed data integration and weighted nearest neighbor (WNN) clustering using RNA and protein expression. Differential gene expression (DGE) was subsequently used to identify 11 distinct cell types (**Fig. 1D, Supplementary Fig. 1D-F**) with canonical gene (**Fig. 1E, Supplementary Fig. 2A**) and protein (**Fig. 1F, Supplementary Fig. 2B**) signatures.

To assess cell specific transcriptional changes in heart failure (HF), we performed pseudobulk DGE analysis at the patient level and observed strong differences between donor and HF (acute MI, ICM, NICM) within fibroblasts, endothelium, smooth muscle cells (SMCs) and pericytes, and myeloid cells (**Fig. 1G**). We also observed marked differences in cell type composition across conditions (**Fig. 1H, Supplementary Fig. 1F**). Increased proportions of myeloid and T-cells and reduced endothelial cell abundance was observed following acute MI with the greatest expansion in myeloid cells occurring within the first week. Accumulation of fibroblasts was seen across pathological conditions beginning 1 week after acute MI. We calculated cell densities using a Gaussian kernel in the WNN UMAP space and uncovered prominent shifts in fibroblasts, myeloid cells, and endothelium phenotypes between donor, acute MI, and chronic HF (ICM, NICM) conditions.

### Expansion of pathologic fibroblast subsets in acute MI and chronic heart failure

Unbiased clustering, harmony integration to correct for batch effects, and subsequent DGE identified 13 distinct fibroblast cell states (**Fig. 2A**) marked by classical fibroblast markers (F1, ground state), *ACTA2/TAGLN* (F2, myofibroblasts), *CCL2/THBS1* (F3), *APOE/AGT* (F4), *DLK1/GPX3* (F5), *PTGDS/GPC3* (F6), *APOD/IGFBP5* (F7), *GDF15/ATF5* (F8), *FAP/POSTN* (F9), *ISG15/MX1* (F10), *PLA2G2A* (F11), *PCOLCE2/MFAP5* (F12), and *PRG4/CXCL14* (F13) (**Fig. 2B**). Construction of gene set signatures for each fibroblast cell state confirmed clear transcriptional delineation between fibroblast states (**Supplementary Fig. 3A-B**). Gene Ontology (GO) analysis based on differentially expressed marker genes identified unique pathway enrichment across states (**Supplementary Fig. 4A**). Spatial transcriptomics data from human heart indicated that fibroblast populations were located within defined perivascular and interstitial spatial niches (**Supplementary Fig. 4B – E**)^25^.

**Figure 2.**
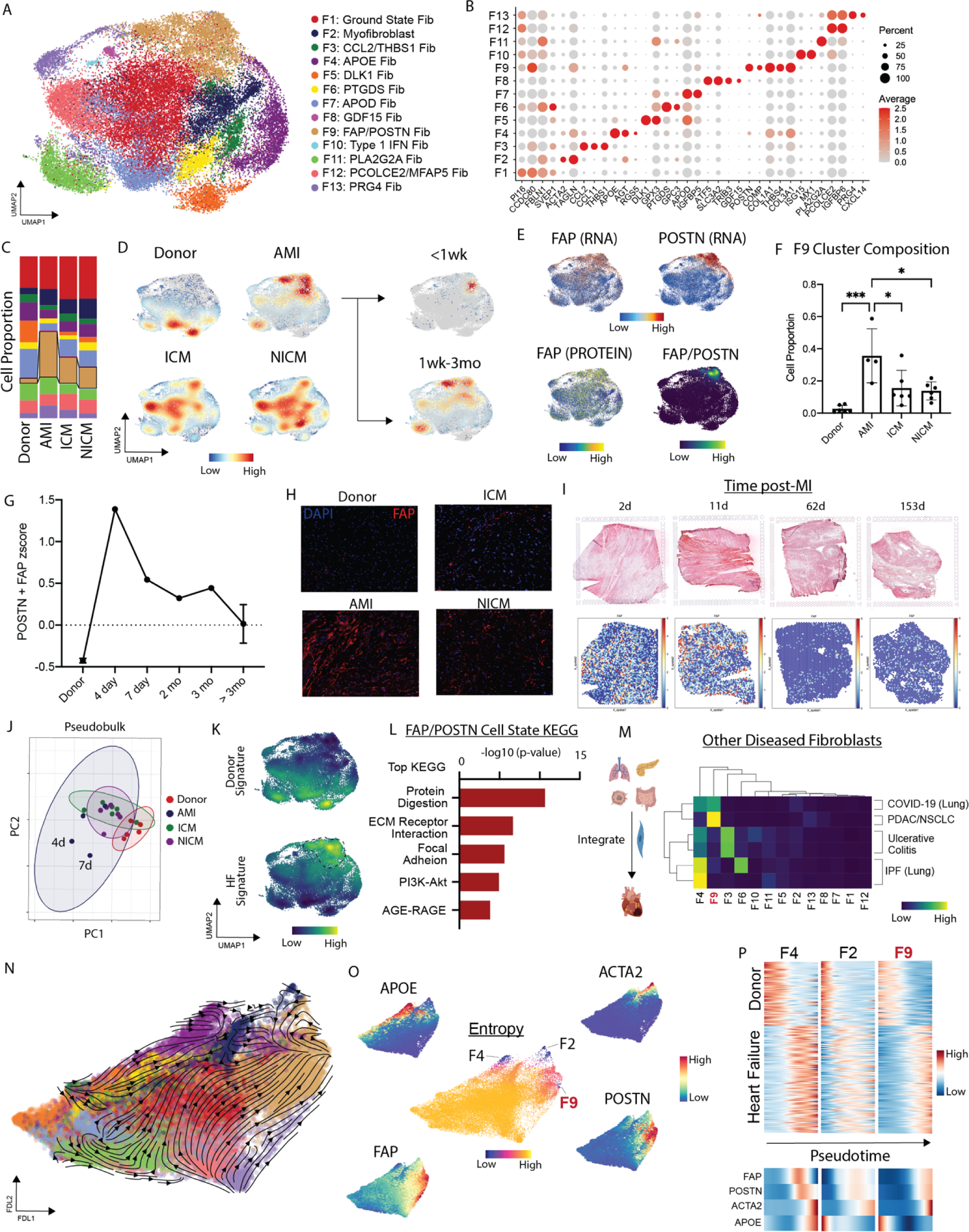
FAP fibroblasts expand post myocardial infarction. (A) UMAP embedding of fibroblast cell states. (B) Dot Plot of marker genes for fibroblast cell states. (C) Fibroblast cell state composition across four groups. (D) Gaussian kernel density estimation of cells across four groups (left) and split by MI time in acute MI patients. (E) FAP RNA (top left), POSTN RNA (top right), FAP protein (bottom left), and FAP/POSTN joint density (bottom right). (F) F9: FAP/POSTN cluster composition across four groups. One-way ANOVA test with multiple comparisons: donor vs acute MI (***P = 0.0002), acute MI vs ICM (*P = 0.0207), and acute MI vs NICM (*P = 0.0115). (G) FAP/POSTN z-score in donor, acute MI, and ICM split by time from MI. (H) Immunofluorescence of FAP in donor, acute MI (4 days post-MI), ICM, and NICM left ventricle myocardium. (I) FAP expression from spatial transcriptomics sections (bottom) with corresponding H&E (top) at different times post-MI. (J) Pseudobulk principal component analysis plot colored by HF category. (K) Gene set signature kernel density embedding plot for donor (top) and HF (bottom) – genes are derived from pseudobulk differential expression analysis between donor and HF. Statistically significant genes (adjusted p-value < 0.05, log2FC > 0.58, and base mean expression > 500). (L) Top KEGG pathways for FAP/POSTN cell state. (M) Pearson correlation coefficient between human heart fibroblasts and those from other disease contexts. (N) scvelo analysis in force directed layout embedding and (O) Palantir derived entropy with terminal cell states noted and marker genes for terminal states in addition to FAP plotted in FDL embedding. (P) Heatmap of pseudobulk differentially expressed genes between donor and HF and terminal state marker genes over pseudotime.

Fibroblast cell states differed across donor, acute MI, and chronic heart failure conditions. Donor hearts contained the greatest number of F4 (*APOE/AGT*) and F5 (*DLK1/GPX3*) fibroblasts. We used an existing Visium spatial transcriptomics dataset (Methods) from healthy LV tissue and found strongest enrichment of F4, F5, and F6 fibroblast gene signatures – F9 and F13 showed almost no signal while F2 was enriched in the perivascular space (**Supplementary Fig. 4B, C**). Acute MI and chronic heart failure patients contained the greatest proportion of F9 (*FAP/POSTN*) and F2 (myofibroblasts) (**Fig. 2C-D**). F9 fibroblasts were strongly enriched for *FAP* at the RNA and cell surface protein levels co-localizing with *POSTN* (**Fig. 2E**). Differential expression analysis from the cell surface protein data identified additional markers for the F2 (F11R) and F9 (THY1, CD276, LAMP1, CD63) states (**Supplementary Fig. 3C-D**). In addition to differing transcriptional signatures, pathway enrichment, and protein markers, F2 and F9 fibroblasts resided within distinct regions of the myocardium (**Supplementary Fig. 4D**). F2 fibroblasts were localized in perivascular regions, while F9 fibroblasts were associated with fibrosis and collagen expression. Collectively, these data highlight that myofibroblasts and FAP/POSTN fibroblasts represent distinct disease associated fibroblast populations. Given the fibrosis associated transcriptional signature of F9 fibroblasts and their localization within the infarct and fibrotic regions of the myocardium, we chose to focus our analysis on this population.

Quantification of the proportion of F9 fibroblasts in individual patients demonstrated consistent increases in acute MI, ICM, and NICM compared to donors (**Fig. 2F**). Temporally, the *FAP/POSTN* transcriptional signature peaked early after MI, declined over a 3 month period, and remained chronically elevated thereafter (**Fig. 2G**). Immunostaining for FAP in LV sections showed no expression in donors, robust expression following acute MI, and moderate expression in ICM and NICM (**Fig. 2H**). To further validate our findings, we utilized spatial transcriptomics data from human MI at days 2, 11, 62, and 153 post-MI. Within this dataset, *FAP* expression was most prominent acutely after MI and diminished at later time points **(Fig. 2I**). We next performed pseudobulk DGE analysis between donor and diseased fibroblasts (**Fig. 2J**) and created gene set signatures for donor and diseased fibroblasts using genes with p-adjusted <0.05, baseMeanExpression >500, and log2FC >0.58. The diseased signature localized specifically within F9 fibroblasts, while the donor signature localized within F5, F7, and F11 fibroblasts (**Fig. 2J-K**). KEGG pathway analysis using F9 marker genes (p-adjusted <0.05 and log2FC >0.58) revealed enrichment for ECM remodeling, cell-cell adhesion, protein digestion, PI3K-Akt signaling, and AGE-RAGE pathways (**Fig. 2L**).

To assess how cardiac fibroblasts the emerge in acute MI and chronic heart failure compare to fibroblasts from other disease contexts, we utilized a published single-cell dataset that examined diseased fibroblasts in 3 human tissues: pancreas (cancer), intestine (ulcerative colitis), and lung (COVID-19, non-small cell lung cancer, idiopathic pulmonary fibrosis)^26^. We re-normalized the published data with SCTransform, integrated it with our human cardiac fibroblasts dataset, and performed clustering using RNA information (**Supplementary Fig. 5A-C**). To assess similarity among integrated fibroblast clusters, we calculated Pearson correlation coefficients between pairwise cell states and found that most cardiac fibroblasts differed from fibroblasts found within other tissues and diseases. F9 fibroblasts were most similar to cancer fibroblasts and F4 fibroblasts were similar to IPF fibroblasts (**Fig. 2M**).

To map fibroblast trajectories in acute MI and heart failure, we undertook two approaches. First, we used scVelo with dynamical modeling to infer directionality in fibroblast cell state differentiation. Projection of the velocity vector field in a force directed layout suggested covergence toward the F2, F4, and F9 fibroblast states (**Fig. 2N**). Next, we used Palantir to predict fibroblast differentiation endpoints, which also identified F2, F4, and F9 as possible terminal states with low entry values across a range of simulation parameters (**Fig. 2O**). To glean how fibroblasts might evolve over the time course of disease, we plotted donor and diseased fibroblast signatures (**Fig. 2K**) versus pseudotime along each predicted lineage (F2, F4, F9) (**Fig. 2P**). The donor fibroblast signature was most enriched early in pseudotime, while the diseased signature increased over pseudotime in each lineage (**Fig. 2O**). However, *FAP* expression increases and then falls along the F2/F4 lineage while it monotonically increases along the F9 lineage. Notably, the markers of each population increase over pseudotime in a lineage specific manner (**Fig. 2P**).

### Mouse cardiac injury models recapitulate human cardiac fibroblast diversity

To address whether available experimental systems are suitable to study cardiac fibroblasts including the F9 fibroblast, we examined several mouse cardiac injury models and human fibroblast culture systems. First, we sought to address how well a traditional mouse left anterior descending artery (LAD) ligation model mimics human disease using previously published scRNAseq data obtained from mouse hearts at day 0, 1, 3, 5, 7, 14, and 28 following MI^11^. Raw data was reprocessed using the same computational pipeline employed in our CITE-seq human data to ensure consistency. We first mapped the mouse MI dataset onto the global human CITE-seq reference and detected strong mapping across all populations (**Supplementary Fig. 6A-B**). Consistent with our human data, we observed increase myeloid infiltration on days 1-7 post-MI (**Supplementary Fig. 6C**) and phenotypic shifts within fibroblast, myeloid, and endothelial cell populations (**Supplementary Fig. 6D-E**). We then mapped mouse fibroblasts onto our human CITE-seq fibroblasts using label-transfer to impute cell state annotation. Mapping scores were strong across all most cell states except for F5, F6, and F7 fibroblasts, which were poorly represented in the mouse data (**Fig. 3A**, **Supplementary Fig. 7A**). When split by time point, there was marked expansion of F9 fibroblasts that peaked on day 7 post-MI and remained elevated compared to sham on d28 post-MI (**Fig. 3B, Supplementary Fig. 7B**).

**Figure 3.**
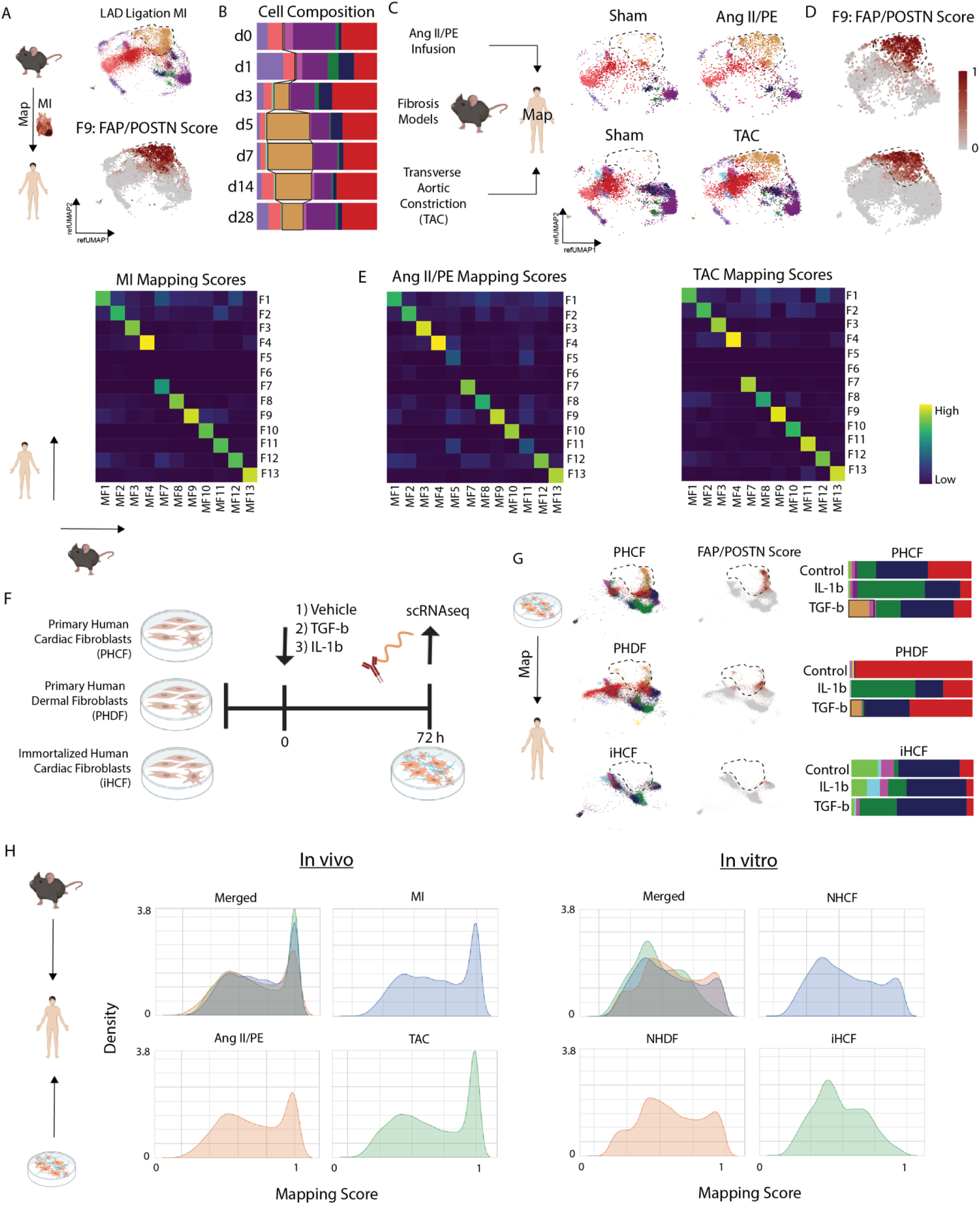
Comparison of *in vivo* and *in vitro* models to study fibroblast fate specification post-MI. (A) Reference mapped mouse MI fibroblasts onto human space with label transfer (top) and prediction score for cells annotated in POSTN/FAP cluster. (B) Cell composition from label-transferred cell states post-MI. (C) Reference mapped mouse fibrosis model fibroblasts from Ang II/PE and TAC split by sham and injury. (D) Prediction score for cells annotated in POSTN/FAP cluster from fibrosis models. (E) Heatmap of average cell prediction score of human fibroblast cell states (y-axis) in annotated mouse fibroblasts (x-axis) in MI, Ang II/PE, and TAC. (F) Experimental set-up of *in vitro* model. (G) Reference mapped *in vitro* cell lines post-stimuli (control, TGF-b, and IL-1b) onto human cardiac fibroblasts and corresponding cell composition. (H) Density plot of maximum prediction score for each cell from *in vivo* and *in vitro* systems merged and split by *in vivo* or *in vitro* condition.

Next, we assessed other mouse cardiac fibrosis models. We infused angiotensin II and phenylephrine (AngII/PE) via implanted osmotic minipump and performed scRNA-seq at day 28 (**Supplementary Fig. 8A**). After applying QC filters (**Supplementary Fig. 8B**), data normalization, clustering, and cell type annotation (**Supplementary Fig. 8C-D**), we selected the fibroblasts and performed label transfer onto our human cardiac fibroblast CITE-seq dataset (**Fig. 3C, Supplementary Fig. 8E**). Consistent with mouse MI, we recovered most fibroblast cell states except for F5 and F6. We observed expansion of F9 fibroblasts, which were absent in the sham control cohort (**Fig. 3C-E**). We then examined another published scRNAseq dataset from hearts that underwent transverse aortic constriction (TAC)^27^. We reprocessed the data and performed label transfer to map mouse TAC fibroblasts onto fibroblasts from our human CITE-seq dataset (**Supplementary Fig. 9A-B**). Consistent with mouse MI and Ang II/PE infusion models, we found strong mapping scores for most fibroblast populations except for F5 and F6. Robust expansion of F9 fibroblasts relative to sham was also seen (**Fig. 3C-E, Supplementary Fig. 9A**). Intriguingly, we observed that JQ1 treatment, which resulted in improved LV systolic function and abrogation of fibrosis, was associated with the loss of F9 fibroblasts. Following JQ1 withdrawal, F9 fibroblasts re-emerge suggesting that this is a reversible effect (**Supplementary Fig. 9B**).

To examine whether commonly used human cultured fibroblast systems model human cardiac fibroblasts, we tested three different fibroblast cell preparations: primary human cardiac fibroblasts (PHCF), primary human dermal fibroblasts (PHDF), and immortalized human cardiac fibroblasts (iHCF). Each fibroblast cell line was treated with vehicle, TGF-β, or IL-1β for 72 hours and scRNAseq performed. Multiple biological and technical replicates were included, individually hash tagged, and pooled to minimize any potential batch effects (**Fig. 3F**). After applying QC filters and data normalization, the data was integrated with harmony (**Supplementary Fig. 10, 11A-B**). To compare human fibroblast preparations to human cardiac fibroblasts, we performed label transfer to map the normalized data onto our human CITE-seq fibroblast dataset and imputed cell annotations. In general, PHCF and PHDF recovered a broader diversity of fibroblast cell states compared to iHCF. However, all cultured cell lines displayed a limited ability to recapitulate fibroblast populations that reside within the human heart. TGF-β treatment led to a modest expansion in F9 fibroblasts in the PHCF and PHDF models, however these cells displayed suboptimal mapping to the human CITE-seq dataset (**Fig. 3G**). To directly compare the ability of mouse cardiac fibroblasts and human cell culture preparations to recapitulate the diversity of human cardiac fibroblasts, we constructed density histograms of the human CITE-seq mapping scores. Fibroblasts recovered from the mouse cardiac injury models showed superior mapping scores relative to the human cell culture preparations (**Fig. 3H, Supplementary Fig. 11C-D**). These data indicate that mouse cardiac injury models are a reasonable experimental approach to explore mechanisms involved in cardiac fibrosis including *FAP/POSTN* fibroblasts.

### Inflammatory signaling within spatially defined cell neighborhoods drives cardiac fibrosis

To identify cell-cell signaling events that might participate in cardiac fibrosis, we used NicheNet to probe ligand-receptor signaling events enriched in diseased conditions relative to donors. We focused on signals received by fibroblasts from any cell type and identified TGF-β and IL-1β as the strongest predicted signals (**Fig. 4A**). We chose to focus on IL-1β signaling given the availability of FDA approved therapeutics and potential clinical impact. *IL-1β* was specifically expressed in myeloid cells (**Fig. 4B**). Within the myeloid cell compartment, *IL-1β* expression was observed in *CCR2^+^* monocytes, macrophages, and classic dendritic cells (DC2s) (**Fig. 4C, Supplementary Fig. 12A-F**). As an orthogonal approach, we plotted *IL-1β* expression in spatial transcriptomics data obtained from a patient following acute MI (day 2) and observed strong expression in the ischemic zone overlapping with the *CCR2^+^* monocytes, macrophages, and cDC2s (**Fig. 4C, Supplementary Fig. 12G, H**). IL-1β expression was conserved in mouse CCR2^+^ monocytes and macrophages in the AngII/PE infusion model (**Fig. 4D**).

**Figure 4.**
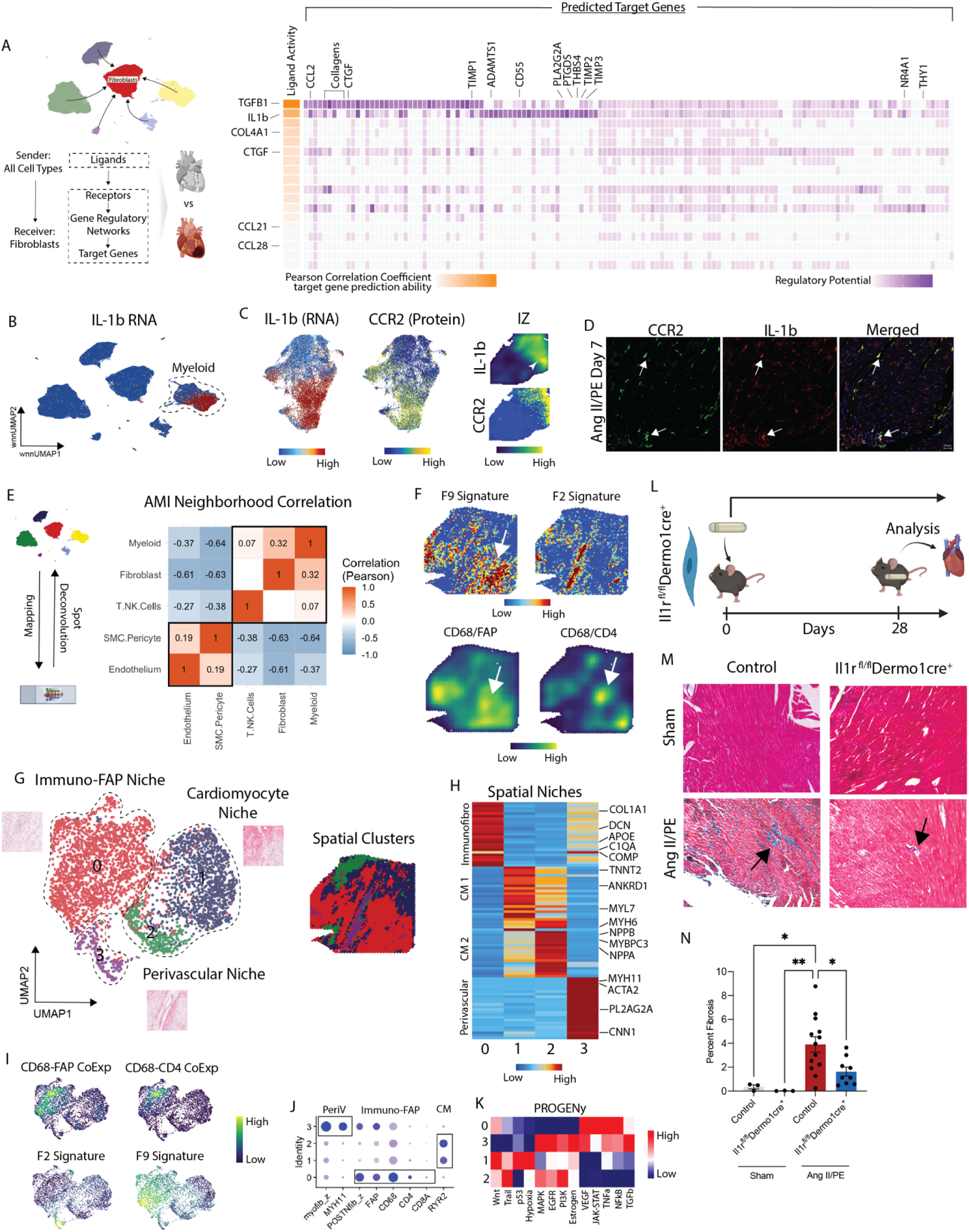
CCR2 macrophages co-localize with and signal to fibroblasts along the IL-1b axis. (A) NicheNet derived predictions of enriched signaling from other cell types to fibroblasts in HF (acute MI, ICM, NICM) relative to donor. (B) IL-1b RNA expression in global CITE-seq UMAP embedding. (C) IL-1b RNA expression in myeloid subsets (left) with corresponding CCR2 protein expression (middle), and IL-1b expression density and imputed CCR2 protein expression in an acute MI IZ spatial transcriptomic sample. (D) Immunofluorescence of IL-1b (red) in CCR2 GFP (green) mice at day 7 post Ang II/PE mini-pump implantation. (E) SPOTlight derived cell-cell proportion correlation in acute MI patient. (F) z-score for marker genes in F9 (*POSTN, COMP, FAP, COL1A1, THBS4, COL3A1*) and F2 (*ACTA2, TAGN*) (top) and CD68/CD4 and CD68/FAP joint density embedding plot (bottom). (G) UMAP embedding plot of spatial clusters with three distinct niches and corresponding spatial location of clusters. (H) Heatmap of top differentially expressed genes across spatial niches. (I) CD68/CD4, CD68/FAP (top) and F2 and F9 fibroblast gene signature (bottom) joint density embedding plot in UMAP embedding. (J) Dot plot of marker genes and gene set scores for spatial niches. (K) PROGENy pathway analysis for spatial clusters in (G). (L) Study design for Il-1r^fl/fl^Dermo1cre mice with Ang II/PE minipump implantation and quantification of fibrosis at day 28.

To determine whether myeloid cells are spatially localized within regions that contain *FAP/POSTN* fibroblasts in the diseased human heart, we carried out the following analysis. We performed snRNA-seq in 7 acute MI and ICM LV specimens and integrated the data with our previously published heart failure atlas (consisting of donor and NICM hearts) to generate a comprehensive single nuclei map of the healthy, injured, and failing human heart than can be used to annotate the spatial transcriptomic data (**Supplementary Fig. 13A**). We identified 13 distinct major cell types as using DGE testing. We then selected representative donor, acute MI, and ICM LV spatial transcriptomic datasets for further analysis. Consistent with our other analyses *FAP* expression was increased early after MI within the infarct area and remained elevated in ICM (**Supplementary Fig. 13B**). To localize cell types within space, we first plotted gene signatures for major cell populations and found consistent overlap with anatomic structures (cardiomyocytes, blood vessels, infarct) identified from the H&E images (**Supplementary Fig. 13C**). To capture cell types which co-localize with fibroblasts in acute MI, we used SPOTlight to deconvolute each spot into its relative cell composition using our reference. After deconvolution, we calculated cell-cell correlations and found fibroblasts most strongly co-localize with myeloid cells and T-cells forming an immune-fibroblast niche. SMCs/pericytes and endothelial cells co-localize in distinct perivascular regions (**Fig. 4E**). We applied this approach to multiple donor, acute MI, and ICM patient samples and identified similar niches (**Supplementary Fig. 14-15**). We then plotted gene signatures for fibroblast states that expanded in disease (F2, F9) and found that F9 fibroblasts co-localized with immune-fibroblast niche (*CD68/FAP/CD4*), while F2 fibroblasts were found within the perivascular niche (**Fig. 4F**).

To examine transcriptional signatures represented in defined niches, we clustered spatial spots within a UMAP embedding and identified 4 spatial clusters (**Fig. 4G**). Cluster 0 was strongly enriched for fibroblast and immune cell markers, clusters 2 and 3 expressed cardiomyocyte markers, and cluster 3 was enriched with SMC and pericyte markers (**Fig. 4H**). Cluster 0 co-localized with regions in space enriched for the F9 gene set score derived from the human CITE-seq dataset (**Fig. 4J**). Co-localization of *CD68/FAP* and *CD68/CD4* was also present in the immune-fibroblast niche. Myofibroblasts (F2) were not present within this niche and instead localized to perivascular regions (**Fig. 4I**). We then used PROGENy to understand what pathways are enriched in the spatial clusters and found enrichment for NFkB signaling (downstream mediator of IL-1β) in the immune-fibroblast niche. TGF-β signaling was most represented in the perivascular niche (**Fig. 4K**). To validate our findings in a broader array of patients, we integrated spatial transcriptomic data across 28 samples in UMAP space from donor, acute MI, and ICM patients. We found high *FAP* and *POSTN* expression in neighborhoods containing expression of macrophage markers (**Supplementary Fig. 16A-C**). These regions contained the F9 fibroblast signature and were highest in the infarct zone (IZ) followed by the fibrotic zone (FZ) and lowest in donors and the remote zone (RZ) (**Supplementary Fig. 16D**).

To determine whether IL-1b signaling to fibroblasts is necessary for cardiac fibrosis, we generate mice that lack the IL-1 receptor (IL1R) in fibroblasts (*Il1r^flox/flox^Dermo1-cre* mice). Control and *Il1r^flox/flox^Dermo1-cre* mice either underwent sham surgery or received Ang II/PE eluting osmotic minipumps. Fibrosis was quantified 4 weeks later by trichrome staining (**Fig. 4L**). Ang II/PE infusion resulted in increased interstitial fibrosis in control mice. Consistent with the conclusion that IL-1 signaling to fibroblasts promotes cardiac fibrosis, we observed reduced trichrome staining in AngII/PE treated *Il1r^flox/flox^Dermo1-cre* mice compared to controls (**Fig. 4M, N**).

### IL-1**β** inhibition impedes FAP fibroblast fate specification and maturation

To dissect the therapeutic impact of IL-1β inhibition on fibroblast cell state transitions, we implanted mice with Ang II/PE osmotic minipumps treated animals with an IL-1β neutralizing antibody (anti-IL-1β mAb) or isotype control every 3 days (**Fig. 5A**). Sham mice served as a negative control. To characterize *FAP* fibroblast fate specification we developed a flow cytometry assay to detect FAP cell surface expression on mouse cardiac fibroblasts (**Supplementary Figure 17**). Mice were harvested after 7 days on AngII/PE treatment and hearts enzymatically digested for flow cytometry. Quantification of flow cytometry data revealed an increase in the proportion of FAP^+^ fibroblasts in AngII/PE treated animals that received isotype antibodies compared to corresponding controls. Importantly, we observed a reduction in FAP^+^ fibroblasts in AngII/PE treated mice that received anti-IL-1β mAb compared to AngII/PE treated mice that received isotype antibody (**Fig. 5B**) demonstrating that endogenous IL-1β signaling regulates FAP^+^ fibroblast specification.

**Figure 5.**
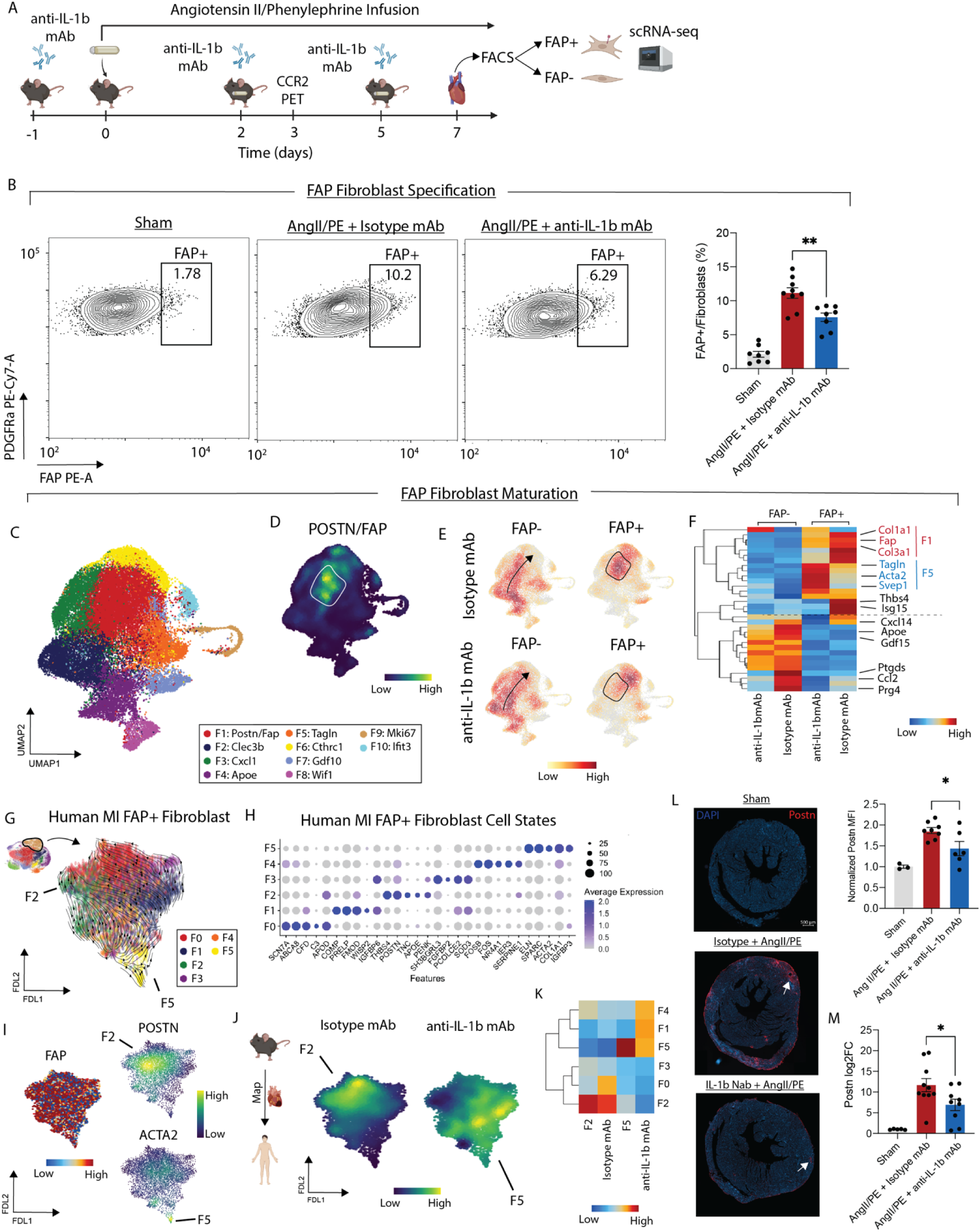
(A) Experimental workflow for cardiac injury experiment with treatment of an IL-1b neutralizing antibody or isotype control. (B) Flow cytometry gating of PDPN vs FAP in a representative sham, Ang II/PE + Isotype, and Ang II/PE + anti-IL-1b mAb mouse heart (left) and FAP+ fibroblasts/total fibroblasts quantified (right). Unpaired t-test and **P = 0.0031 (isotype vs anti-IL-1b mAb). (C) Integrated UMAP for isotype and anti-IL-1b mAb treated mice FAP+/FAP- fibroblasts with annotated clusters. (D) Density plot of Fap/Postn co-localization with area of maximal expression highlighted. (E) Gaussian kernel density plots of 4 conditions in integrated UMAP embedding. (F) Heatmap of key fibroblast cell state genes grouped by four conditions with rows clustered by similarity. (G) scVelo of human FAP+ fibroblasts in a FDL embedding space colored by re-clustered data. (H) Top marker genes for FAP+ human CITE-seq re-clustered data from (F). (I) FAP, POSTN, and ACTA2 expression in human FAP+ fibroblasts in FDL embedding. (J) Mapping mouse differentially expressed signature between FAP+ isotype and anti-IL-1b mAb treated mice. (K) Heatmap of gene set signature for F2 (*THBS4, POSTN, TNC, APOE,* and *PENK*), isotype treated, F5 (*ACTA2* and *TAGLN*), and anti- IL-1b mAb treated grouped by FAP+ human fibroblast cell state. (L) IF of *POSTN* in a representative sham, Ang II/PE + Isotype, and Ang II/PE + anti-IL-1b mAb heart (left) with quantification (right). Unpaired t-test and *P = 0.0243. (M) RT-PCR of Postn in mouse hearts split by 3 conditions. Unpaired t-test and *P = 0.0392.

To delineate whether IL-1β signaling contributes to FAP^+^ fibroblast maturation, we sorted FAP^+^ and FAP^-^ fibroblasts from the hearts of mice that were treated with AngII/PE for 7 days and received either isotype or anti-IL-1β mAb. Sorted fibroblasts were then subjected to scRNAseq (**Fig. 5A**). Following application of QC filters and data normalization (**Supplementary Fig. 18A-B)**, cells were clustered into 10 transcriptionally distinct fibroblast cell states (**Fig. 5C, Supplementary Fig. 18B**). Density plot of *Fap/Postn* co-expression showed strong enrichment in clusters F1 and F6 (**Fig. 5D**). GO Biological Pathway analysis identified enrichment for fibrosis associated pathways including ECM organization, integrin signaling, and collagen fibrin organization in mouse FAP^+^ fibroblast states (**Supplementary Fig. 18C**).

To unravel phenotypic shifts among fibroblasts following anti-IL-1β mAb treatment, we constructed Gaussian kernel density embedding plots. As anticipated, significantly different cell distributions were found in the FAP^+^ and FAP^-^ groups with enrichment of cells in the F1 and F6 clusters in the FAP^+^ fibroblast group. Mice that received anti-IL-1β mAb displayed reductions in FAP^+^ fibroblasts that localized to cluster F1 compared to corresponding isotype control (**Fig. 5E**). We detected differences in gene expression in both the FAP^+^ and FAP^-^ fibroblasts following anti-IL-1β mAb treatment (**Supplementary Fig. 18D**). Within the FAP^+^ population, isotype treated animals displayed increased expression of *Fap, Thbs4, Isg15, Col1a1, and Cola3.* Conversely, IL-1β mAb treatment increased the expression of myofibroblast markers including (*Tagln*, *Acta2*, and *Svep1*) in FAP^+^ fibroblasts (**Fig. 5F**). These data suggest that IL-1β mAb treatment may impact the differentiation trajectory of FAP^+^ fibroblasts. GO biological pathway analysis further suggested that anti-IL-1β mAb treatment impacted genes involved in ECM organization, T cell migration, cytokine signaling, and chemokine production (**Supplementary Fig. 19D**). FAP^-^ fibroblasts showed reduced expression of *ApoE*, *Gdf15*, *Ptgds*, *Ccl2*, and *Prg4* following IL-1β mAb treatment (**Fig. 5F**).

To examine the differentiation trajectory of FAP^+^ fibroblasts, we sub-clustered human FAP^+^ fibroblasts (F9 cell state) from our human CITE-seq dataset at finer resolution. DEG analysis showed distinct cell states within the FAP^+^ populations (**Fig. 5G-H**). All cells expressing *FAP* and only a subset expressing *POSTN* or *ACTA2* (**Fig. 5I**). To predict differentiated cell states, we used scVelo and observed convergence within the *POSTN* fibroblast (F2) and *ACTA2* myofibroblast (F5) clusters (**Fig. 5G**). Next, we mapped the mouse FAP^+^ fibroblasts onto the human FAP^+^ fibroblast FDL trajectory and plotted gene signatures enriched in following either isotype or anti-IL-1β mAb treatment. We observed a strong separation triggered by anti-IL-1β mAb treatment (**Fig. 5J**). To interpret the effect of anti-IL-1β mAb treatment within the context of the human FAP^+^ fibroblast differentiation, we created a heatmap of isotype enriched, anti-IL-1b mAb enriched, and human *POSTN* (F2) signatures grouped by human FAP^+^ fibroblast state. We found that the isotype signature corresponds with the human F2 (*POSTN*) signature, while the anti-IL-1b mAb signature is enriched for alternative human fibroblasts states including myofibroblasts (F1, F4, F5) (**Fig. 5K**). These data further suggest that IL-1b mAb treatment impedes the maturation of FAP^+^ fibroblasts into *POSTN* fibroblasts.

To experimentally validate these findings, we performed immunostaining for POSTN in sham hearts and hearts treated with AngII/PE for 7 days. Compared to sham controls, AngII/PE infusion increased the abundance of POSTN^+^ cells. Consistent with our scRNAseq data, we observed a reduction in POSTN^+^ cells in AngII/PE treated hearts from mice that received anti-IL-1β mAb compared to isotype control (**Fig. 5L**). Similarly, quantitative PCR revealed upregulation of *Postn* expression in myocardial tissue obtained from AngII/PE treated mice compared to sham controls. Treatment with anti-IL-1β mAb reduced *Postn* expression induced by AngII/PE infusion within the heart relative to isotype control (**Fig. 5M**).

## Discussion

The advent of single cell multi-omic technologies has afforded the opportunity for the high-resolution detection of novel cell states from healthy and diseased human tissue including the heart^19, 20, 22, 23, 25, 28^. These human cell atlases have provided new insights into the cellular landscape of HF and identified unexpected cell diversity within the stromal compartment. However, the functional roles of these novel cell states and how they might interact during disease pathogenesis remains to be defined. Among stromal cell types, fibroblasts are of particular interest given their role in tissue fibrosis, arrhythmias, and heart failure^16, 29, 30^. Little is known regarding the mechanisms and signaling mechanisms that orchestrate the specification of pathogenic fibroblasts in the context of human MI and HF. Herein, we utilize CITE-seq and spatial transcriptomics to dissect fibroblast cell states that emerge following acute MI and in chronic HF and identify a fibroblast trajectory marked by *FAP* and *POSTN* expression that are governed by inflammatory cytokines and contribute to cardiac fibrosis.

Unbiased clustering revealed an expansion of fibroblasts expressing *FAP* that peaked early after MI and remained elevated in chronic HF. We used IF and spatial transcriptomics in acute MI hearts^25^ to validate expression of FAP at the RNA and protein levels. Recent studies have utilized chimeric antigen receptor (CAR) T cells to target FAP^+^ fibroblasts in a mouse model of cardiac injury and shown improved cardiac function and diminished fibrosis^9, 10^. This points to a potential pathogenic role of FAP^+^ fibroblasts in adverse LV remodeling. Transcriptionally, cardiac FAP^+^ fibroblasts resemble cancer associated fibroblasts, which also express *FAP*^3126^. Computational lineage analysis predicted that *FAP* expression marks a fibroblast trajectory that transitions into populations including POSTN^+^ matrifibrocytes^32^ and myofibroblasts. Unraveling the signals that drive the differentiation and maturation of FAP+ fibroblasts may identify new therapeutic targets to limit fibrosis in the injured and failing heart.

A fundamental question that continues to challenge the field is selection of appropriate model systems to investigate human cardiac fibroblasts. To address this issue, we leveraged publically available and new scRNAseq datasets to examine the suitability of mouse cardiac injury models (MI, Ang II/PE infusion, TAC) and human cultured fibroblast platforms. Surprisingly, we found that each of the mouse models contained many of the fibroblast populations found in the human heart and recapitulated expansion of *Fap* and *Postn* expressing fibroblasts following injury. In contrast, primary human cardiac fibroblasts, primary human dermal fibroblasts, and immortalized human cardiac fibroblasts were relatively poor models of human cardiac fibroblast diversity under resting or stimulated (TGF-β^20, 33, 34^, IL-1β^35^) conditions. These findings emphasize the relevance of *in vivo* mouse models in studying cardiac fibrosis and for therapeutic discovery.

Little is understood regarding the signaling mechanisms that underlie the specification and differentiation of pathological fibroblasts in the heart. Using cell-cell interaction analysis, we identified enriched for IL-1β and TGF-β signaling in fibroblasts in HF relative to donor control conditions. While numerous studies have implicated a role for TGF-β signaling in maladaptive cardiac remodeling and scar formation in HF^20, 33^, the potential for inflammatory signals to drive fibroblast activation and cardiac fibrosis is less explored. In support of this concept, a large clinical trial targeting the IL-1β pathway with the neutralizing antibody, canakinumab, led to a reduction in ischemic cardiac events^36^. We show that IL-1β is selectively expressed by CCR2^+^ monocytes and macrophages in the human heart consistent with our prior mouse studies^37, 38^. Using spot deconvolution and Pearson correlation analysis in spatial transcriptomic data from human hearts, we identified spatial niches of macrophages and T-cells that strongly co-localize with fibroblasts in acute MI. This niche robustly expressed *FAP*, extracellular matrix components associated with fibrosis, and displayed an inflammatory signature. We validated the existence of this immune-fibroblast neighborhood in a larger dataset of 28 patients and observed strong macrophage-fibroblast co-localization in pro-inflammatory niches. Together, these findings suggest that FAP^+^ fibroblasts reside within a spatially defined cell neighborhood that contains inflammatory macrophages and that these cell types may communicate via IL-β signaling.

To verify the importance of IL-1 signaling, we generated mice that lack the IL-1 receptor in fibroblasts and demonstrated a specific requirement for IL-1 signaling to fibroblasts in cardiac fibrosis. Next, we sought to explore the mechanistic implications of inhibiting IL-1β signaling as a therapeutic. We found that anti-IL-1 β mAb treatment abrogated the emergence of FAP^+^ fibroblasts. scRNAseq of cardiac fibroblasts in this model uncovered an unexpected effect on the maturation of the *Fap* fibroblast trajectory. We observed that FAP^+^ fibroblasts differentiated into POSTN^+^ fibroblasts in isotype control hearts. Strikingly, mice treated with the anti-IL-1 β mAb displayed a differentiation trajectory away from POSTN^+^ fibroblasts and towards the myofibroblast lineage. These findings were validated by immunostaining and highlight a central role of IL-1β signaling in pathogenic fibroblast fate specification. They also suggest that IL-1β targeted therapies may not interfere with structural stability of the heart as myofibroblasts are preserved in this context.

Our study is not without limitations. First, we are not powered to assess the effects of demographics (age, sex, ethnicity) on the cell state diversity post-MI. We included patients of diverse ethnicities, sex, and broad age ranges to remain inclusive in the generation of the human dataset. Second, we have a limited cohort size in the AMI group. Given the challenge acquiring fresh acutely infarcted cardiac tissue, we validated our CITE-seq findings in a broad array of patients through multiple assays such as IF and spatial transcriptomics.

In conclusion, we utilize multi-omic sequencing to characterize the inflammatory fibrosis axis in human myocardial infarction and HF. We dissect myeloid-fibroblast crosstalk using spatial transcriptomics, interactome analysis, and *in vivo* perturbation to show that IL-1 β modulates fibroblast specification, maturation, and cardiac fibrosis. Collectively, our findings highlight a promising role for immunomodulators in targeting cardiac fibrosis. These findings have broad implications for future therapeutic discovery in the emerging discipline of cardio-immunology.

## Acknowledgments

KL is supported by the Washington University in St. Louis Rheumatic Diseases Research Resource-Based Center grant (NIH P30AR073752), the National Institutes of Health [R01 HL138466, R01 HL139714, R01 HL151078, R01 HL161185, R35 HL161185], Leducq Foundation Network (#20CVD02), Burroughs Welcome Fund (1014782), and Children’s Discovery Institute of Washington University and St. Louis Children’s Hospital (CH-II-2015-462, CH-II-2017-628, PM-LI-2019-829), Foundation of Barnes-Jewish Hospital (8038-88), and generous gifts from Washington University School of Medicine. JMA is supported by the American Heart Association Predoctoral Fellowship (826325). PM is supported by American Heart Association Postdoc Fellowship (916955). Figure 1C, 6E, and 7A were created in BioRender.com. We thank the Genome Technology Access Center at the McDonnell Genome Institute at Washington University School of Medicine for help with genomic analysis. The Center is partially supported by NCI Cancer Center Support Grant #P30 CA91842 to the Siteman Cancer Center. This publication is solely the responsibility of the authors and does not necessarily represent the official view of NCRR or NIH.

## Author Contributions

JA and AB isolated cells and made cDNA from human CITE-seq samples. TY made all CITE-seq libraries for sequencing. PM, AB, and AK isolated nuclei for snRNA-seq. JA and XL performed all computational analysis. JA, CK, and SH analyzed spatial transcriptomics data. AF and SS performed human *in vitro* experiments. JA performed all *in vivo* mouse surgeries and downstream experiments. JA, SY, VP, FK, and GF performed immunohistochemistry and analyzed and processed images. VP completed all work associated with the Il1r^fl/fl^ Dermo1cre mice. JA and KL drafted the manuscript. MF contributed to the experimental design, interpretation, and manuscript production. TY, C-ML, SS and BA contributed to the experimental design, data analysis and interpretation as well as manuscript production. KL is responsible for all aspects of this manuscript including experimental design, data analysis, and manuscript production. All authors approved the final version of the manuscript.

## Materials and Methods

### Ethical Approval for Human Specimens

The study is compliant with all relevant ethical regulations and has been approved by the Washington University School of Medicine Institutional Review Board (IRB #201104172). Informed consent was obtained from each patient prior to tissue collection by Washington University School of Medicine and no compensation was provided in exchange for subject participation in the study. All demographic and clinical data has been de-identified and provided in Supplementary Table 1. Patients included in this study span diverse race, age, and sex to provide an inclusive trans-ethic study population.

### Inclusion Criteria

Prior to tissue collection, specific inclusion criteria were employed to ensure well controlled study groups. Any patients with HIV or hepatitis and known genetic cardiomyopathies were excluded from this study. Donor hearts: patients with stable ejection fractions, no know history of cardiac disease and a non-cardiac cause of death/transplant. Acute MI hearts: patients who had a myocardial infarction from coronary artery disease within 3 months from the time of transplant. Ischemic cardiomyopathy: patients who had a myocardial infarction from coronary artery disease greater than 3 months from the time of transplant. Non-ischemic cardiomyopathy: patients with HF independent of ischemic heart disease and no known history of a myocardial infarction.

### Human single cell isolation for CITE-seq

Fresh cardiac left ventricular tissue from ex-planted hearts at the time of transplantation, LVAD cores, or donors declared DCD at Washington University School of Medicine and Mid America Transplant Service were harvested, perfused with cardioplegia, and transported on ice. In donor, ICM, and NICM hearts a section of the LV apex was dissected out; in acute MI hearts a section of the infarct region (identified as a white discoloration) was dissected. Tissues were minced using a razor blade on ice and transferred to a 15 mL conical tube containing 3mL DMEM with 170uL Collagenase IV (250U/mL final concentration), 35uL DNAse1 (60U/mL), and 75uL Hyaluronidase (60U/mL) and incubated at 37 °C for 45 min with agitation. After 45 min, the digestion reaction was quenched with 6 mL of HBB buffer (2% FBS and 0.2% BSA in HBSS), filtered through 40 µm filters into a 50 mL conical tube and transferred back into a 15 mL conical tube to obtain tighter pellets. Samples were then spun down at 4 °C, 1200rpm for 5 min and the supernatant was discarded. Pellet resuspended in 1 mL ACK Lysis buffer (Gibco A10492-01) and incubated at room temperature for 5 min followed by the addition of 9 mL DMEM and centrifugation (4 °C, 5 min, 1200 rpm). Supernatant was discarded and the pellet was resuspended in 2 mL FACS buffer (2% FBS and 2mM EDTA in calcium/magnesium free PBS) and centrifugation was repeated in above conditions and supernatant aspirated. The TotalSeq A 277 panel (BioLegend) antibody cocktail was resuspended in 100 uL of FACS buffer and 1 uL each of custom oligo-labeled FAP (Amgen) and LRRC15 (Amgen) were added. The combined 102 uL were used to resuspend the pellet with the addition of 1 uL of DRAQ5 (Thermo Fisher Scientific, 564907) and incubated on ice for 30 min. Solution was washed 3x with FACS buffer following same centrifugation as above and then resuspended in 500 uL of FACS buffer and 1 uL DAPI (BD Biosciences, 564907) and filtered into filter-top FACS tubes. First singlets were gated and subsequent DRAQ5+/DAPI- events were collected in 300 uL cell resuspension buffer (0.04% BSA in PBS) – collected cells were centrifuged as above and resuspended in collection buffer to a target concentration of 1,000 cells/uL. Cells were counted on a hemocytometer before proceeding with the 10x protocol.

### CITE-seq Library Preparation

Collected cells were processed using the using the single Cell 3’ Kit v 3.1 (10x Genomics PN: 1000268). 10,000 cells were loaded onto ChipG (PN:1000121) for GEM generation. Reverse transcription, barcoding, and complementary DNA amplification of the RNA and ADT tags were performed as recommended in the 3’ v3.1 chromium protocol. Single-cell libraries were prepared using the single Cell 3’ Kit v 3.1 following a modified 3’ v3.1 assay protocol (User Guide CG000206) to concurrently prepare gene expression and TotalSeq A antibody derived tag (ADT) libraries as recommended by BioLegend. 1 ul of 0.2uM ADT Additive Primer (CCTTGGCACCCGAGAATT*C*C) and 15 ul of cDNA Primers (PN: 2000089) were used to amplify cDNA. ADT libraries were amplified with a final concentration of 0.25 uM SI Primer (AATGATACGGCGACCACCGAGATCTACACTCTTTCCCTACACGACGC*T*C) and 0.25 uM TrueSeq Small RNA RPI primer (CAAGCAGAAGACGGCATACGAGAT[6nt index]GTGACTGGAGTTCCTTGGCACCCGAGAATTC*C*A) using 15 cycles. Gene expression libraries were indexed using Single Index Kit T Set A (PN: 2000240). Libraries were sequenced on a NovaSeq 6000 S4 flow cell (Illumina).

### CITE-seq alignment, quality control, and cell type annotation

Raw fastq files were aligned to the human GRCh38 reference genome (v) using CellRanger (10x Genomics, v6.1) with the antibody capture tag for the TotalSeqA 277 + two custom oligo-tagged antibodies. First, protein reads were normalized and de-noised for each sample separately using the dsb package^39^ in R v4 with the isotype controls tag to remove noisy cell-to-cell variation. Subsequent quality control, normalization, dimensional reduction, and clustering was performed in Seurat v4.0. Following normalization, quality control was performed and cells passing the following criteria were kept for downstream processing: 500 < nFeature_RNA < 600 and 1,000 < nCount_RNA < 25,000 and percentage mitochondrial reads < 15%. Raw RNA counts were normalized and scaled using SCTransform^40^ regressing out percent mitochondrial reads and nCount_RNA. Principal component analysis was performed on normalized RNA counts and to determine the number of PCs to use for further processing the following criteria was applied: PCs exhibit cumulative percent > 90% and the percent variation associated with the PCs as < 5%. Weighted nearest neighbor clustering^41^ (WNN) was performed with the significant RNA PCs and dsb normalized proteins (minus isotype controls) directly without PCA as previously outlined with the FindMultiModalNeighbors function in Seurat. Subsequently, a uniform manifold approximation (UMAP) embedding was constructed and FindClusters was used to un-biasedly cluster cells using the SLM modularity optimization algorithm. Clustering was performed for a suite of different resolutions (0.1-0.8 at 0.1 intervals) and differential gene expression using the FindAllMarkers function and a Wilcoxon Rank Sum test with a logFC cut-off of 0.25 and a min.pct cut-off of 0.1. Clusters were annotated using canonical gene and protein markers and subsequent violin plots (RNA) and heatmaps (protein) were created to assess clean separation of clusters into distinct cell types. Previously identified canonical marker genes were also plotted on the UMAP object to further validate cluster annotations. Cell composition in donor, acute MI, ICM, and NICM was calculated as percentages as previously described. Cellular density profiles in the WNN UMAP space were calculated in Scanpy per condition using the tl.embedding_density function. Briefly, a Gaussian density kernel estimation is used to estimate cell density in WNN UMAP space in each condition (donor, acute MI, ICM, and NICM) and density values are scaled between 0-1. Similar approach used for all cell density calculations in manuscript.

### Cell composition and density shift calculations

We used R to compute cell type composition across conditions. To assess shifts in cell density within both the global object and within individual cell types, we converted the .rds object to a .h5ad file format and used scanpy.tl.embedding function which employs a Gaussian kernel density estimation of cell number within the UMAP embedding. Density values are scaled from 0-1 within that category.

### Pseudobulk Differential Gene Expression

We use the DESeq2^42^ package to perform pseudobulk DE analysis. After QC, cells were subsetted for each cell population annotated from the global reference, raw counts were aggregated to the patient level, data normalized using a regularized log transform, and differential expression analysis between donor and all HF (acute MI, ICM, NICM) via DESeq2. Genes were deemed statistically significant if adjusted p-value < 0.05 and absolute(log2FC) > 0.58. For constructing pseudobulk donor/HF gene set scores, we only used statistically significant genes with a mean log base expression of > 500.

### Fibroblast, myeloid, and T cell states analysis

To cluster cell types into distinct cell states, we subsetted the cell type of interest, re-normalized, computed PCAs, harmony integrated, computed UMAPs, and clustered data at a range of resolutions. DE analysis was then used to identify marker genes for each cell state and a dot-plot or heatmap to assess clustering separation. Using the top marker genes we calculated gene set z-scores and plotted them in UMAP space.

### Perturbed Disease Associated Fibroblasts

Perturbated state human fibroblasts were used from (E-MTAB-10324)^26^ processed using the same pipeline as used for human CITE-seq fibroblasts. Seurat was then used to integrate the perturbed state human diseased fibroblasts with human HF CITE-seq fibroblasts. To assess cluster similarity, we used two approaches: (1) built a phylogeny tree using the BuildClusterTree function and (2) computed Pearson correlation coefficients and displayed results as a matrix.

### Pseudotime Trajectory Analysis

Palantir: Palantir was used to perform pseudotime analysis on fibroblasts and macrophage subsets. For the cell type of interest, a normalized count matrix was exported and processed in python via Palantir. A force directed layout was then computed for visualization of trajectories and MAGIC was used to impute data for visualization. For fibroblasts, F1 was used as the starting cell state and for myeloid cells monocytes were used as the starting cells and resident macrophages were excluded from the trajectory analysis. No terminal states were specified. Generalized Additive Models within Palantir was then used to compute gene expression trends along the different fibroblast lineages. Pseudotime, entropy, and FDL embedding values were exported and subsequent plotting for visualization was performed in R/Seurat. scVelo: Using CellRanger aligned BAM files as input, a loom file containing the estimation of spliced and unspliced RNA counts were generated by Velocyto CLI (v0.17.17)^43^. The loom file was further merged with annotated fibroblasts object and pre-processed, RNA velocity was estimated using dynamical or stochastic modeling and the fdl embedding stream graph was generated by scvelo (https://github.com/theislab/scvelo).

### Pathway Analysis and transcription factor enrichment

Statistically significant DE genes were used to perform pathway analysis via EnrichR (https://maayanlab.cloud/Enrichr/). Pathway enrichment values were downloaded as .csv files and plots generated in Prism. GO enrichment analysis to compare cell states was done in R using clusterProlifer^44^ compareClusters function using only statistically significant marker genes. PROGENy^45^ was used for pathway analysis in the spatial transcriptomics dataset.

### In Vitro Fibroblast single-cell experiments

Human ventricular cardiac fibroblasts (NHCF-V, Lonza, CC-2904), adult human dermal fibroblasts (NHDF-Ad, Lonza, CC-2511), and immortalized human cardiac fibroblasts (iHCF, Applied Biological Materials Inc., T0446) were grown in FibroGRO LS medium (Millipore, SCMF002) supplemented with 2% (NHDF-Ad) or 10% (NHCF-V and iHCF) FBS. Cells were maintained in a humidified atmosphere (95% air, 5% CO_2_) at 37°C and passaged every 2-3 days. For cytokine treatment, cells were seeded in 6-well plates and allowed to attach for 24hr. Thereafter, cells were equilibrated for 16 hr in low serum medium (0.2% FBS for NHDF-Ad, and 0.5% for NHCF-V and iHCF), and then treated with 10 ng/mL of recombinant human TGF-b (R&D Systems; 240-B-002) or 10 ng/mL of recombinant human IL-1β (R&D Systems; 201-LB-005) for the indicated times. After trypsinization, cell suspensions were labeled with barcoded lipids from the 3’ CellPlex Kit Set A (10x Genomics PN:1000261) following manufacturer’s instructions (Protocol CG000391). After labeling, technical replicates were pooled and loaded onto a Chromium Controller (10x Genomics) at a concentration to capture 6-10,000 cells. Single-cell libraries were prepared using the Single Cell 3’ Kit v3.1 (10x Genomics PN: 1000268) according to manufacturer’s instructions (User Guide CG000388). Gene expression and cellplex libraries were indexed with Dual Index Kit TT Set A and NN Set A indexes (PN: 1000215, 3000482) respectively. Libraries were pooled and sequenced on a NovaSeq 6000 (Illumina).

### Processing of in vitro single cell data

Raw fastq files were aligned to the human GRCh38 reference genome using CellRanger (10x Genomics, v6.1). In vitro single-cell data was processed using the same pipeline as human CITE-seq data for iHCF, NHDF, and NHCF datasets.

### Reference mapping of in vivo and in vitro data

In vitro: The same mapping pipeline was used for iHCF, NHDF, and NHCF datasets. Briefly, SCTransform normalized data was used, the FindTransferAnchors function was used with a spca reference reduction and 50 components. The anchors were then passed into the MapQuery function to impure cell annotations and project the in vitro data into the human CITE-seq fibroblast umap embedding. Mapping scores for each imputed cell state were then plotted in the umap space and a heatmap of average mapping score for each cell state was constructed.

In vivo: for the day 28 Ang II/PE, TAC, and MI the mouse genes were first converted into the human analogs using biomaRt, the data was subsequently normalized using SCTransform, and normalized data was used to find anchors and then projected onto the human CITE-seq fibroblast UMAP space as described with the in vitro data. Mapping of in vivo and in vitro used the same pipeline and parameters.

### Human single nuclei isolation for snRNA-seq

Cardiac tissues from LVAD cores at the time of LVAD implant (U/RR-pre) and adjacent to core samples at the time of explant (U/RR-post) from paired patient were flash frozen using liquid nitrogen. Identical regions from the apex of LV from explanted donors were used. Single nuclei suspensions were generated as previously described. In brief, flash frozen sections were minced with a razor blade, transferred to a Dounce Homogenizer containing 1 mL of lysis buffer (10 mM Tris-HCl, pH 7.4, 10 mM NaCl, 3LmM MgCl_2_ and 0.1% NP-40 in nuclease-free water) on ice. Samples were homogenized using five strokes, an additional 1 mL of lysis buffer added, and incubated on ice for 15 min. Samples were then filtered with a 40 um filter, which was rinsed with 1 mL of lysis buffer. The mixture was then centrifuged at 500 *g* for 5 min 4 °C, resuspended in 1 mL nuclei wash buffer (2% BSA and 0.2LULul^−1^ RNase inhibitor (Thermo Fisher, cat. no. AM2694) in 1X PBS) and, filtered using a 20 um pluristrainer (Pluriselect, cat. No. SKU43-50020-03). Filtered solution as centrifuged using the above criteria and resuspended in 300 uL Nuclei Wash Buffer and transferred into a 5 mL tube for flow cytometry. Subsequently, 1LuL DRAQ5 (5 mM solution; Thermo Fisher, cat. no. 62251) was added, sample gently vortexed and allowed to incubate for 5 min prior to sorting. DRAQ5^+^Lnuclei were sorted into 300 uL Nuclei Wash Buffer using a BD FACS Melody (BD Biosciences) with a 100 uM nozzle. Sorted nuclei were then centrifuged using the above conditions and resuspended in Nuclei Wash Buffer for a final target concentration of 1,000 nuclei/uL – nuclei were counted on a hemocytometer. Based on the nuclei concentration, 10,000 target nuclei were loaded onto a Chip G for GEM generation using the Chromium Single Cell 5L Reagent v1.1 kit from 10X Genomics. Reverse transcription, barcoding, complementary DNA amplification and purification for library preparation were performed as per the Chromium 5L v1.1 protocol at the McDonnel Genome Institute. Sequencing was performed on a NovaSeq 6000 platform (Illumina) at a target read depth of 100,000 at the McDonnel Genome Institute.

### snRNAseq reference map generation

Nuclei fastq files were aligned to the whole genome pre-mRNA reference generated from the GRCh38 transcriptome (with the intron flag included) using CellRanger v3 (10X Genomics). Nuclei were filtered to include those with 1000 < RNA UMI count < 10,000 and mitochondrial reads < 5%. After initial QC, scrublet was ran on each sample separately in Python with default parameters to score nuclei and nuclei with a doublet score > 0.2 were excluded from downstream analysis. We then leveraged supervised doublet removal as previously described on a per cell type basis before combining all objects. Briefly, clusters were annotated into major cell populations, each major cell type was subsetted, and re-normalized, PCA, UMAP embedding, clustering, and DE analysis performed. Sub-clusters which did not express the gene signature of the cell type or had overlapping genes across different cell types were removed. Post-contamination, all cell type objects were merged to construct a cleaned object. After QC and doublet removal downstream analysis was performed in Seurat v4^46^. The cleaned object was normalized using SCTransform^40^ with regressing out mitochondrial percent and RNA UMI counts. We then computed the principal components and used these to integrate all samples with harmony^47^. Informed by the ElbowPlot, we used 80 components to construct the UMAP embedding, find nearest neighbors, and clustered the data at multiple resolutions. We then used the FindAllMarkers function in Seurat to perform differential expression testing and annotated clusters into distinct cell types based on canonical gene markers. To define cell states within each major cell type, we subsetted the major cell populations, re-normalized, re-clustered, re-computed the UMAP, and annotated cell states. Both the global object and each cell type object was saved and used for downstream analysis and plotting in Seurat. All differential gene expression used to identify cell types or cell states was performed using the normalized assay with the FindAllMarkers function and the Wilcoxon rank sum test with a min.pct = 0.1 and logfc.threshold = 0.25.

### Integration of spatial transcriptomics data

Visium 10x data from 10x Genomics was used in Supplementary Figure 3 (https://www.10xgenomics.com/resources/datasets/human-heart-1-standard-1-1-0) – processed as per Seurat v4. Processed .h5ad objects were acquired for all spatial transcriptomic samples as previously published^48^. Data was re-normalized using SCTransform to keep analysis consistent with the human data processing. The snRNA-seq reference map was used to impute voxel annotations from selected spatial transcriptomic samples – mapping scores and gene signatures were also plotted to validate mapping. SPOTlight^49^ was used for spot deconvolution and Pearson correlation coefficients calculated to identify cells which co-localize in space. To validate findings from isolated patients in a broader array of patients, the above analysis was performed on several samples and an integrated UMAP embedding of spatial spots was constructed for 28 samples using the same analysis pipeline as for snRNA-seq reference map construction.

### Receptor-ligand interaction analysis

Using fibroblasts as a receiver, we first used nichenetr (v1.1.0) default pipeline to explore the top 20 ligands from all other non-fibroblast cell types, that are predicted to orchestrate the highest regulatory potential to regulate downstream target expression in fibroblasts. The regulatory potential from top 20 ligands on their predicted targets in fibroblasts was represented in a heatmap. To further capture the disease altered signaling from myeloid cells to fibroblasts, we extended the analysis to draw information from the differential expression of the ligand-receptor pairs across HF groups (HF vs healthy) by using the differential NicheNet pipeline. Circos plot was created to visualize the interaction between ligands from myeloid cells and receptors from fibroblast cells.

### Mouse strains

All animal studies were performed in compliance with guidelines set forth by the National Institutes of Health Office of Laboratory Animal Welfare and approved by the Washington University institutional animal care and use committee. All mouse strains used in this study (WT, CCR2 GFP) are on the C57BL/6 background. Il1r^fl/fl^ mice were crossed to the Dermo1 cre to generate Il1r^fl/fl^Dermo1cre^+^ strain. Il1r^fl/fl^Dermo1cre^-^ mice were used as the control.

### Animal experiments

8–10-week-old male mice were implanted with osmotic minipumps (Alzet model 2001) and constant infusion of angiotensin II (1.5 ug/g/day) and phenylephrine (50 ug/g/day). Mice were treated with either an isotype control or an anti-IL-1b neutralizing antibody (Amgen) at 5 mg/kg via intraperitoneal injection every 3 days starting with a pre-treatment injection one day before osmotic minipump implantation and then at days 2 and 5. Mice at day 7 as described in the relevant sections. For the Il1r^fl/fl^Dermo1cre^+^ Ang II/PE experiment 28 day osmotic minipumps (Alzet model 2004) and mice were sacrificed on day 28 for trichrome staining.

### Fibroblast flow cytometry and sorting for single cell RNA seq analysis

After ice cold PBS perfusion, heart tissue from mice were minced with a razor blade on ice and transferred to a 15 mL conical tube containing 3 mL DMEM with 170 uL Collagenase IV (250 U/mL final concentration), 35 uL DNAse1 (60 U/mL), and 75 uL Hyaluronidase (60 U/mL) and incubated at 37 °C for 45 min with agitation. After 45 min, the digestion reaction was quenched with 6 mL of HBB buffer (2% FBS and 0.2% BSA in HBSS), filtered through 40 um filters into a 50 mL conical tube and transferred back into a 15 mL conical tube to obtain tighter pellets. Samples were then spun down at 4 °C, 1200 rpm for 5 min and the supernatant was discarded. Pellet resuspended in 1 mL ACK Lysis buffer (Gibco A10492-01) and incubated at room temperature for 5 min followed by the addition of 9 mL DMEM and centrifugation (4 °C, 5 min, 1200 rpm). Supernatant was discarded and the pellet was resuspended in 2 mL FACS buffer (2% FBS and 2 mM EDTA in calcium/magnesium free PBS) and centrifugation was repeated in above conditions and supernatant aspirated. Samples were then incubated in Pdpn (BioLegend, 127423, clone 8.1.1), Cd140a (Biolegend, 135911, clone APA5), Cd31 (Biolegend, 102409, clone 390), Fap (Creative Biolabs, HPAB-0171-CN-PE), and Cd45 (Biolegend, 103132, clone 30-F11) with a 1:200 dilution for 30 min. A no Fap negative control was used which contained all antibodies except Fap. Solution was washed 3X with FACS buffer following same centrifugation as above and then resuspended in 500 uL of FACS buffer and 1 uL DAPI (BD Biosciences, 564907) and filtered into filter-top FACS tubes. Gating was performed as follows: Pdpn+, singlets, Cd45-/Cd31-, Pdpn+/PDgfra+, and Fap vs Pdgfra gate was used to look at Fap+ fibroblasts. The no Fap antibody negative control was used to draw the Fap gate.

For the fibroblast scRNAseq experiment, 4 isotype Ang II/PE and 4 anti-IL-1 mAb + Ang II/PE mice FAP+ and FAP- fibroblasts were sorted into 300 uL cell resuspension buffer (0.04% BSA in PBS). Collected cells were centrifuged as above, pooled across mice, and resuspended in collection buffer to a target concentration of 1,000 cells/uL. Cells were counted on a hemocytometer before proceeding with the 10x protocol. cDNA construction and library preparation were performed for 4 libraries (Isotype + Ang II/PE FAP+, Isotype + Ang II/PE FAP-, anti-IL-1b mAb + Ang II/PE FAP+, and anti-IL-1b mAb + Ang II/PE FAP-. cDNA synthesis, library construction, and sequencing were performed using the same protocol as the human CITE-seq data.

### Histology and Immunofluorescence

Flash frozen LV samples were fixed for 24Lhr at 4 °C in 10% neutral buffered formalin, washed in 1X PBS, and embedded in paraffin. Paraffin-embedded sections were cut at an 8Lum thickness using a microtome. Slides were baked at 65 °C for 1 hr, deparaffinized with serial xylene washes, and rehydrated with ethanol. Then slides underwent methanol treatment (10% MeOH + 3% H_2_O_2_) for 20 min at RT followed by 3X TBS-T washes (5 min each). Antigen retrieval was performed using the AR6 buffer (Ankoya Biosciences, AR600250ML) for 15 min in the microwave and then cooled to room temperature. Tissue sections were marked with a hydrophobic pen and blocked in 10% BSA in TBS-T for 30 min at RT. Slides were then stained with the primary antibody diluted in 10% BSA in TBS-T overnight at 4 °C: FAP (Abcam, ab207178, 1:250) and POSTN (Abcam, ab215199, 1:250). Next, the primary antibody was detected using Opal Polymer HRP Ms + Rb (PerkinElmer Opal Multicolor IHC system). The PerkinElmer Opal Multicolor IHC system was utilized to visualize antibody staining per manufacturer protocol. Slides were imaged using a Zeiss Axioscan Z1 automated slide scanner. Image processing was performed using Zen Blue and Zen Black (Zeiss). For POSTN quantification, the whole heart was circled, and an MFI calculated in sham, isotype + Ang II/PE and anti-IL-1b mAb + Ang II/PE representative sections. For the Il1r^fl/fl^Dermo1cre^+^ mice 28-day Ang II/PE infusion, mice were sacrificed on day 28 and FFPE sections prepared as above. Trichrome staining was performed using Gomori’s Trichrome Stain Kit (ThermoScientific, 87020) and subsequently percent fibrosis was quantified in serial cross-sections.

For the CCR2 GFP mice, frozen sections were prepared using the following: hearts perfused with cold PBS, placed in 4% PFA overnight at 4 °C, washed with PBS 3X, infiltrated with 30% sucrose overnight at 4 °C, and embedded in OCT and stored in a −80 °C freezer. Prior to staining, 10 um sections were cut using a Leica Cryostat, washed with PBS, incubated with 0.25% Triton-X in PBS at RT for 5 min, blocked in 5% BSA/PBS for 1hr at RT, stained with primary antibody: anti-GFP (Abcam, ab13970) and IL-1b (R&D systems, AF-401-NA, 1:50) co-stained with an appropriate secondary antibody, and mounted with DAPI fluoroshield (R&D, F6057). Slides were then imaged on a Zeiss confocal microscopy and representative images included.

### RT-PCR

RNA was extracted from flash frozen mouse apex using the Qiagen RNeasy Plus Mini Kit. RNA concentration was then measured using a nanodrop spectrophotometer and cDNA synthesis was performed using the HighCapacity RNA to cDNA synthesis kit (Applied Biosystems). Quantitative real time PCR reactions were prepared with sequence-specific primers with PowerUP™ Syber Green Master mix (Thermo Fisher Scientific) in a 20 uL volume. Real time PCR was performed using QuantStudio3 (Thermo Fisher Scientific). mRNA expression was normalized to 36B4. The Postn primer was purchased from ADT (Fwd: 5’-GGTGCCCTAGAAAGGATCATGG-3’; Rev: 5’-CAGAGCACTGGAGGGTATTTAG-3’).

## Data Availability

Raw sequencing files can be found on the Gene Expression Omnibus (). Processed data is available upon request from the authors.

## Code Availability

Scripts used for analysis in this manuscript can be found at ().

---

**Supplementary Figure 1.**
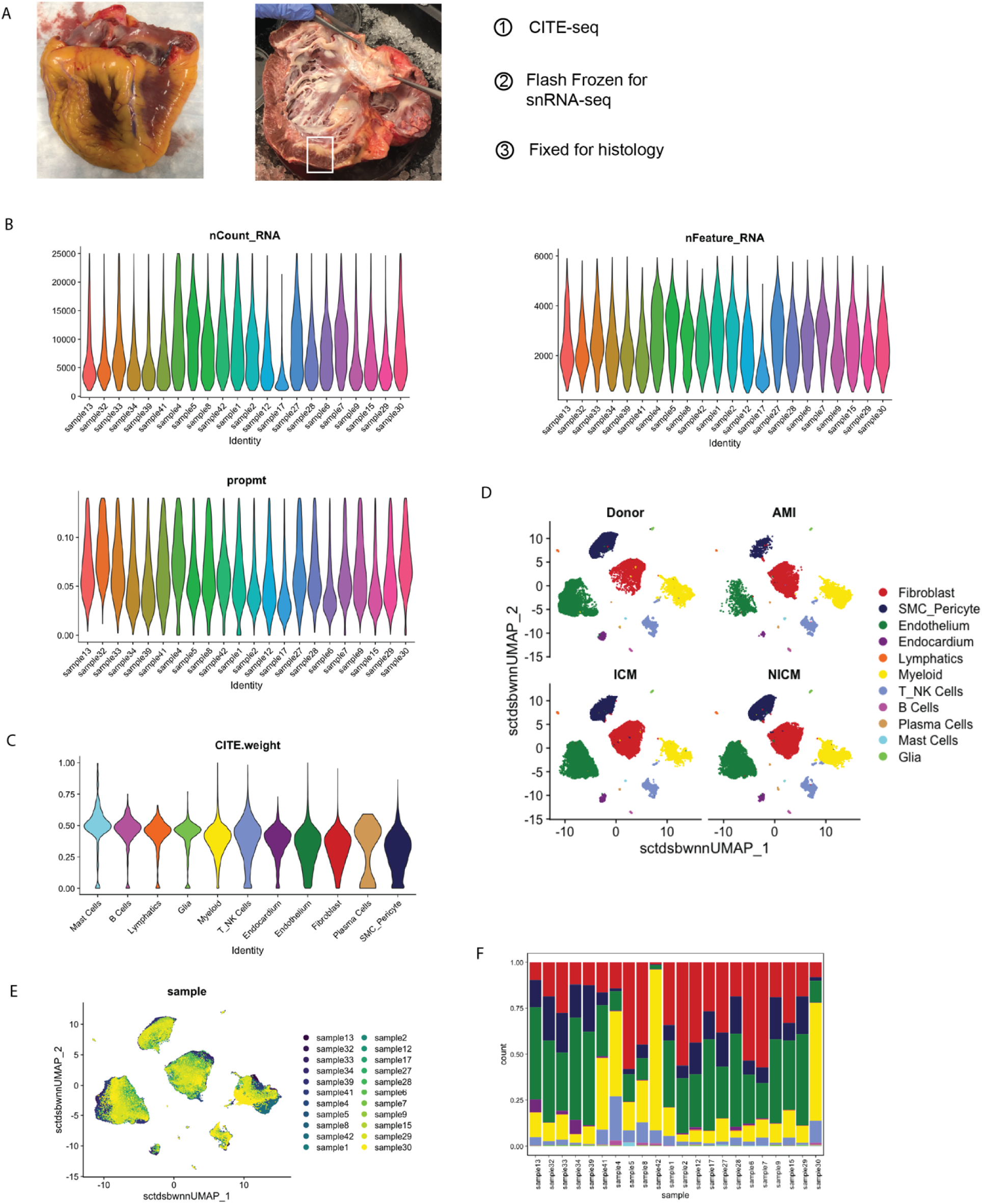
(A) Explanted heart and dissected region boxed used for CITE-seq, snRNA-seq, and histological analysis. (B) QC metrics post filtering. (C) CITE-seq protein assay weights used for WNN clustering. (D) Global integrated UMAP split by 4 conditions. (E) Integrated global UMAP colored by patient sample. (F) Cell type composition split by each sample.

**Supplementary Figure 2.**
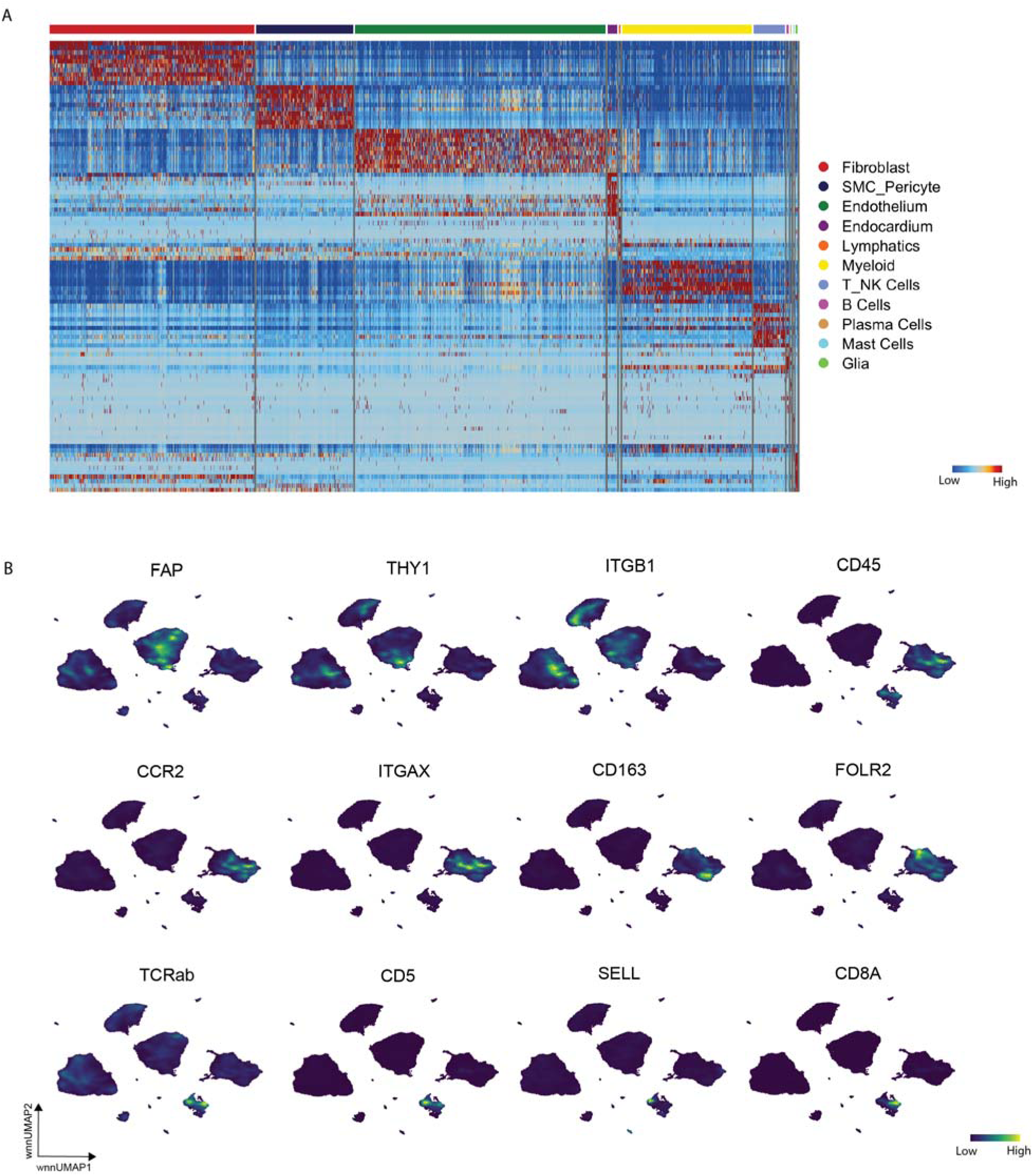
(A) Heatmap of top marker genes for each cell type in global UMAP. (B) Protein expression as a density plot for top CITE-seq protein markers in different cell types.

**Supplementary Figure 3.**
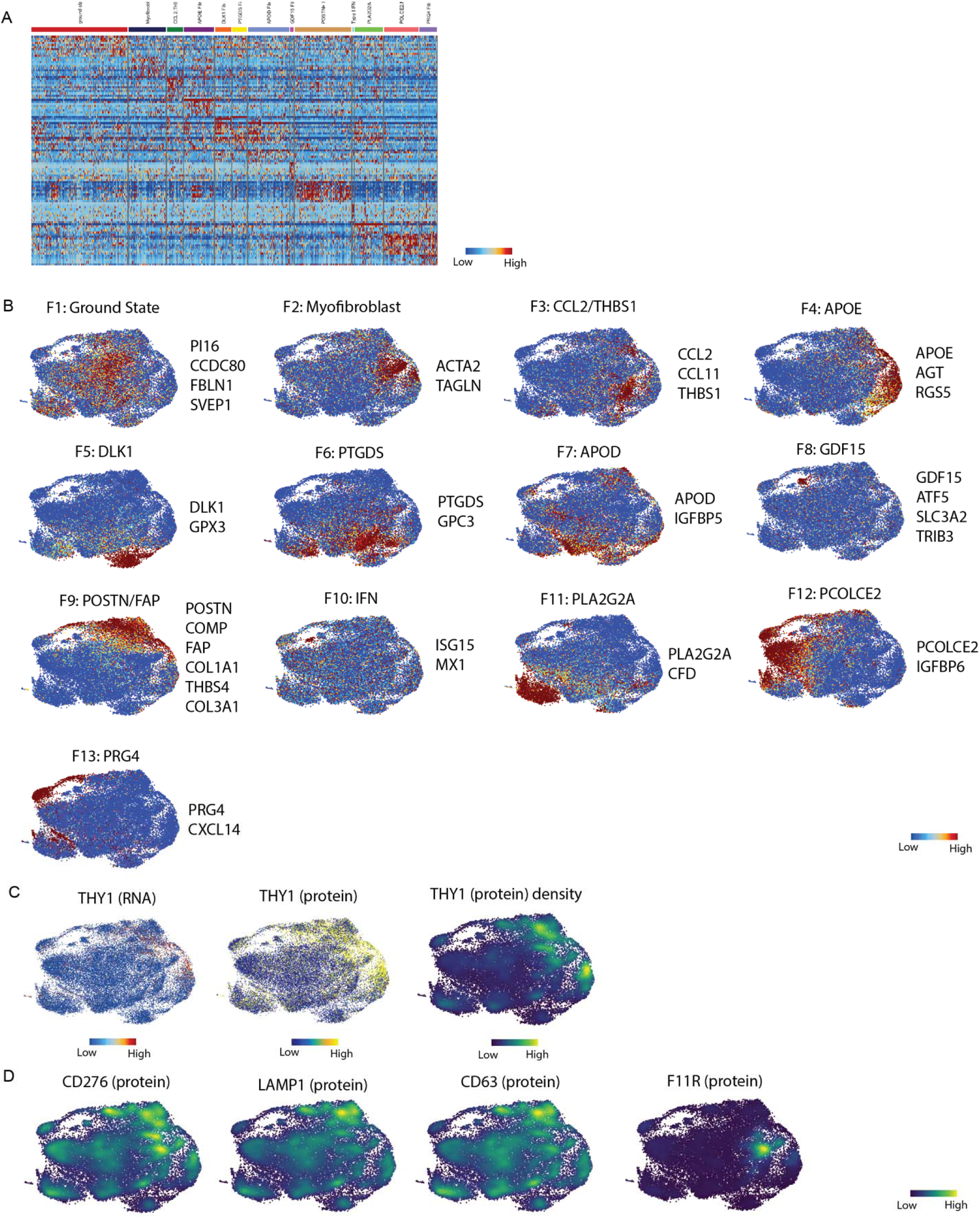
(A) Heatmap of top marker genes for fibroblast cell states. (B) Gene set z-scores for fibroblast cell states plotted in UMAP embedding. (C) THY1 RNA, protein, and protein density plot in fibroblast UMAP space. (D) Density plots for differentially expressed protein markers in fibroblast UMAP space.

**Supplementary Figure 4.**
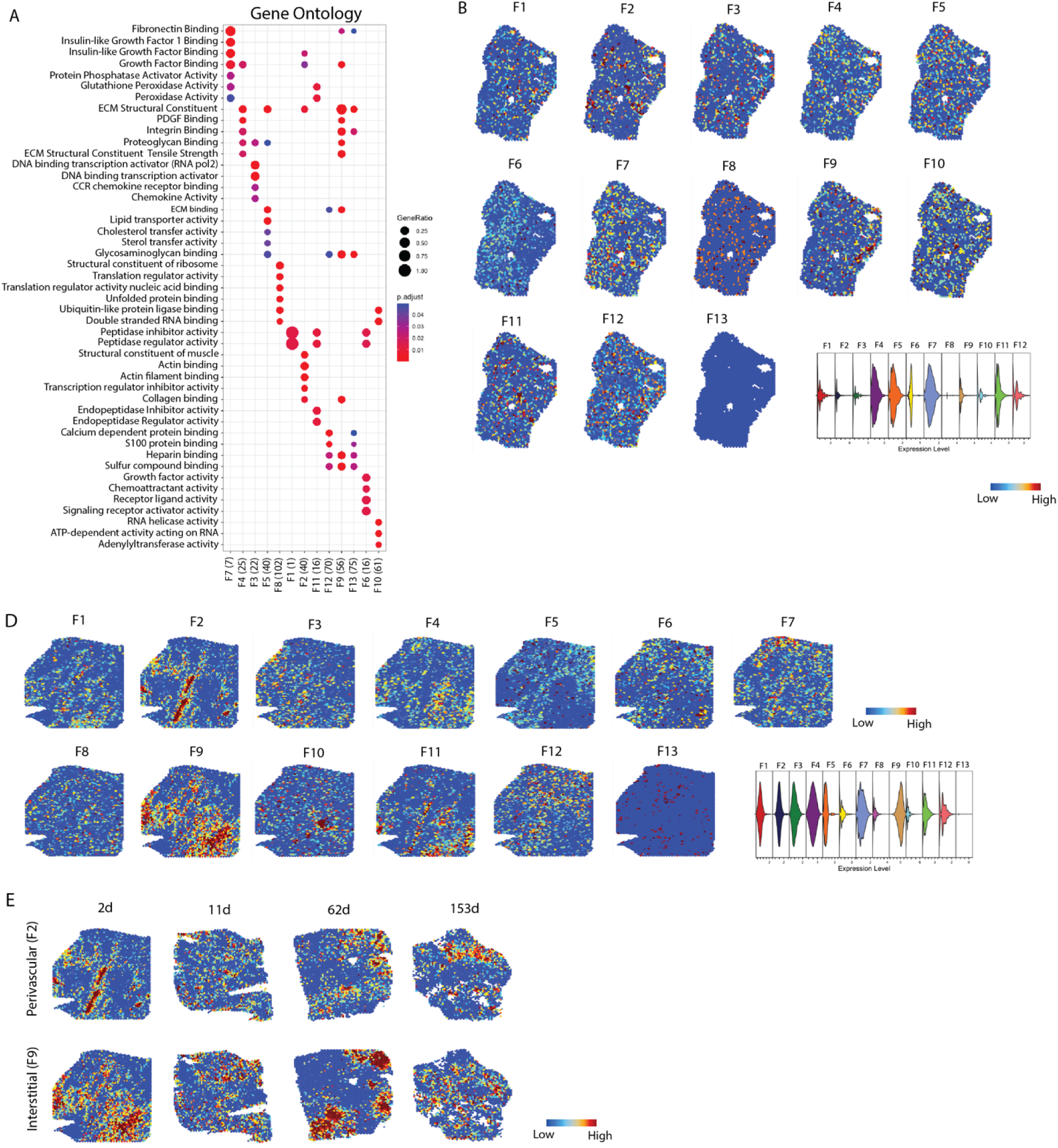
(A) GO analysis from clusterProfiler for fibroblast cell states. Fibroblast cell state gene set scores in a spatial transcriptomics (B) donor and (C) AMI sample with associated violin plot. (D) F2 and F9 gene set score plotted in spatial transcriptomics data across acute MI and ICM sections.

**Supplementary Figure 5.**
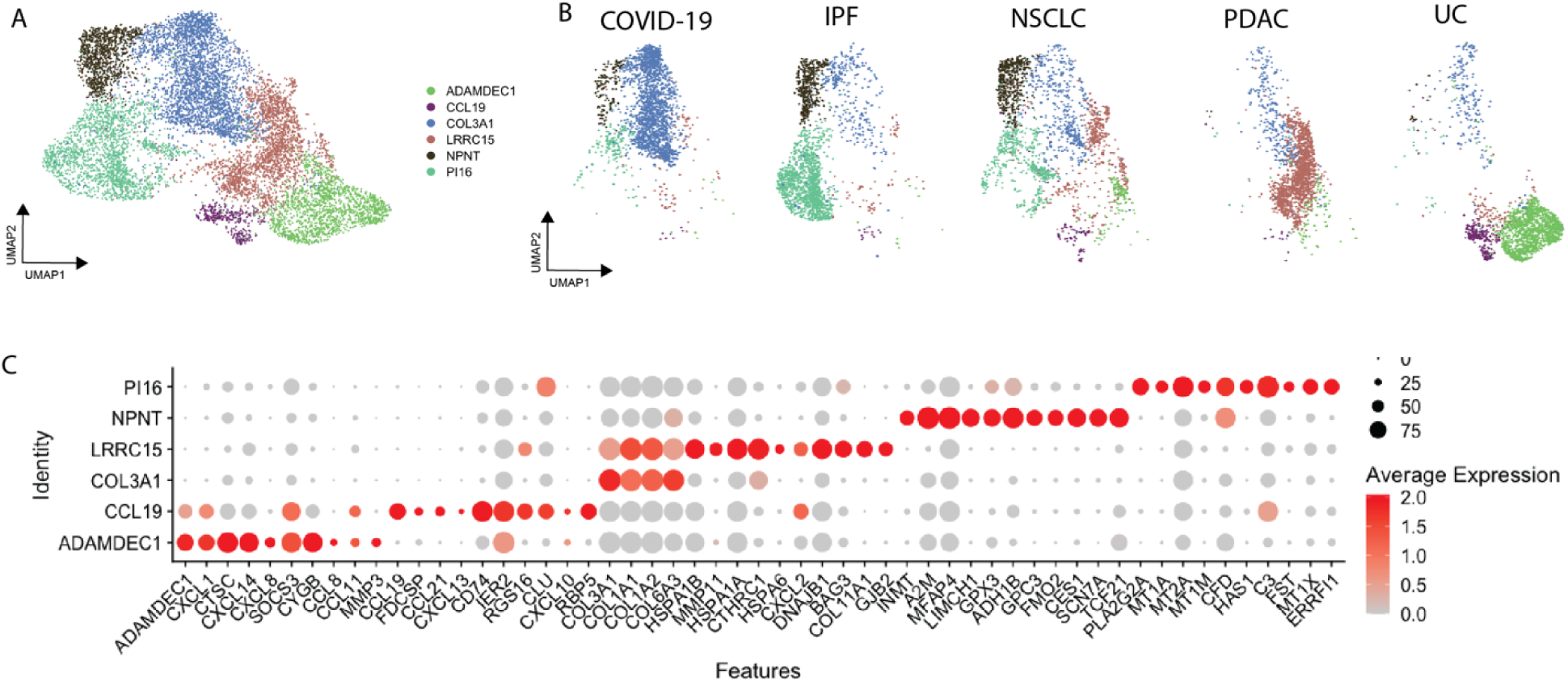
(A) Integrated UMAP of perturbated pathological fibroblasts and (B) split by disease category. (C) Dotplot of marker genes for clusters in (A).

**Supplementary Figure 6.**
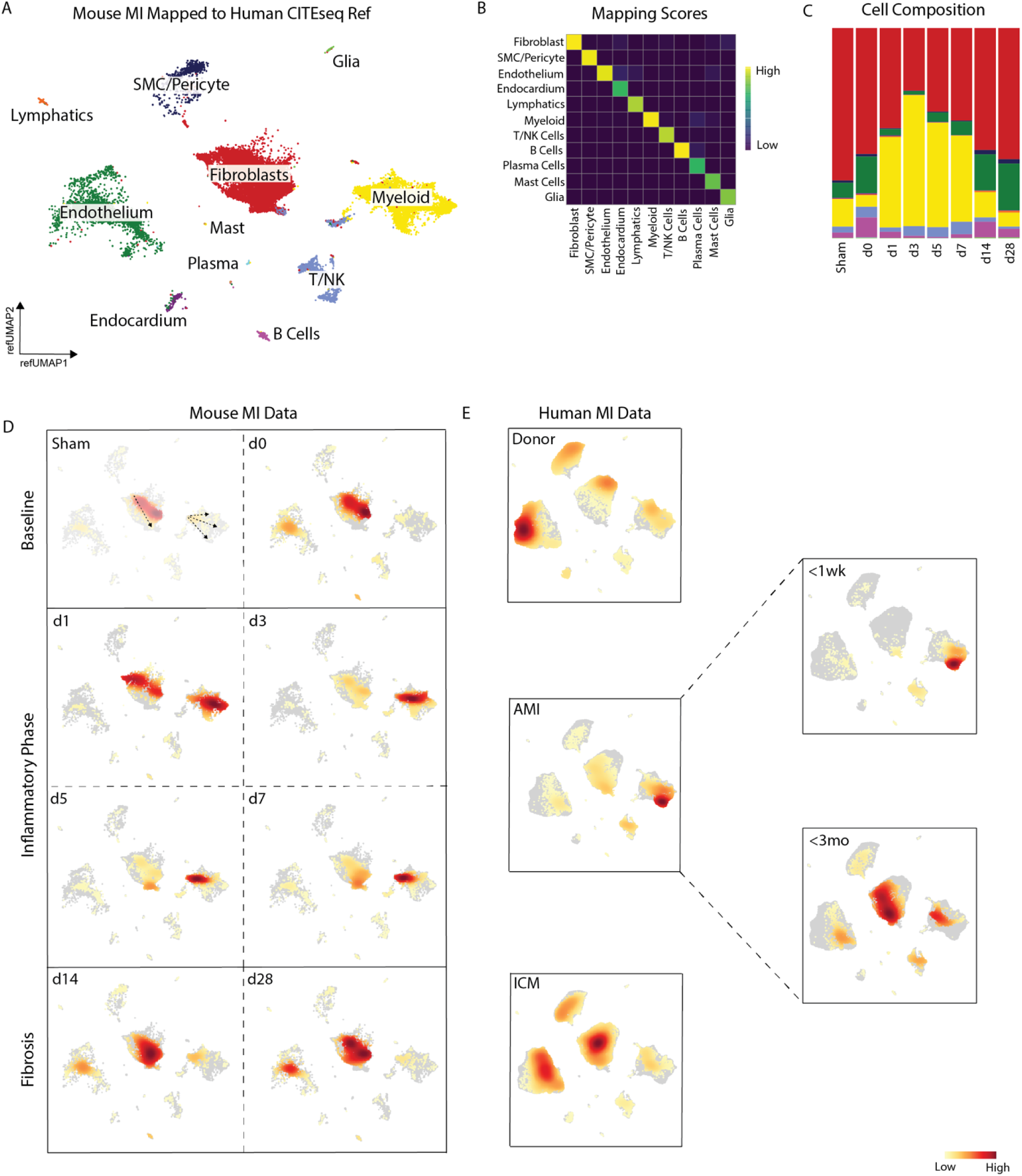
(A) Mouse MI data mapped onto human CITE-seq global object. (B) Seurat reference mapping scores for different cell types. (C) Cell composition from imputed annotations split by time point. (D) Gaussian kernel density embedding plots in mouse MI data split by condition and biological phase and (E) human MI data grouped by MI category.

**Supplementary Figure 7.**
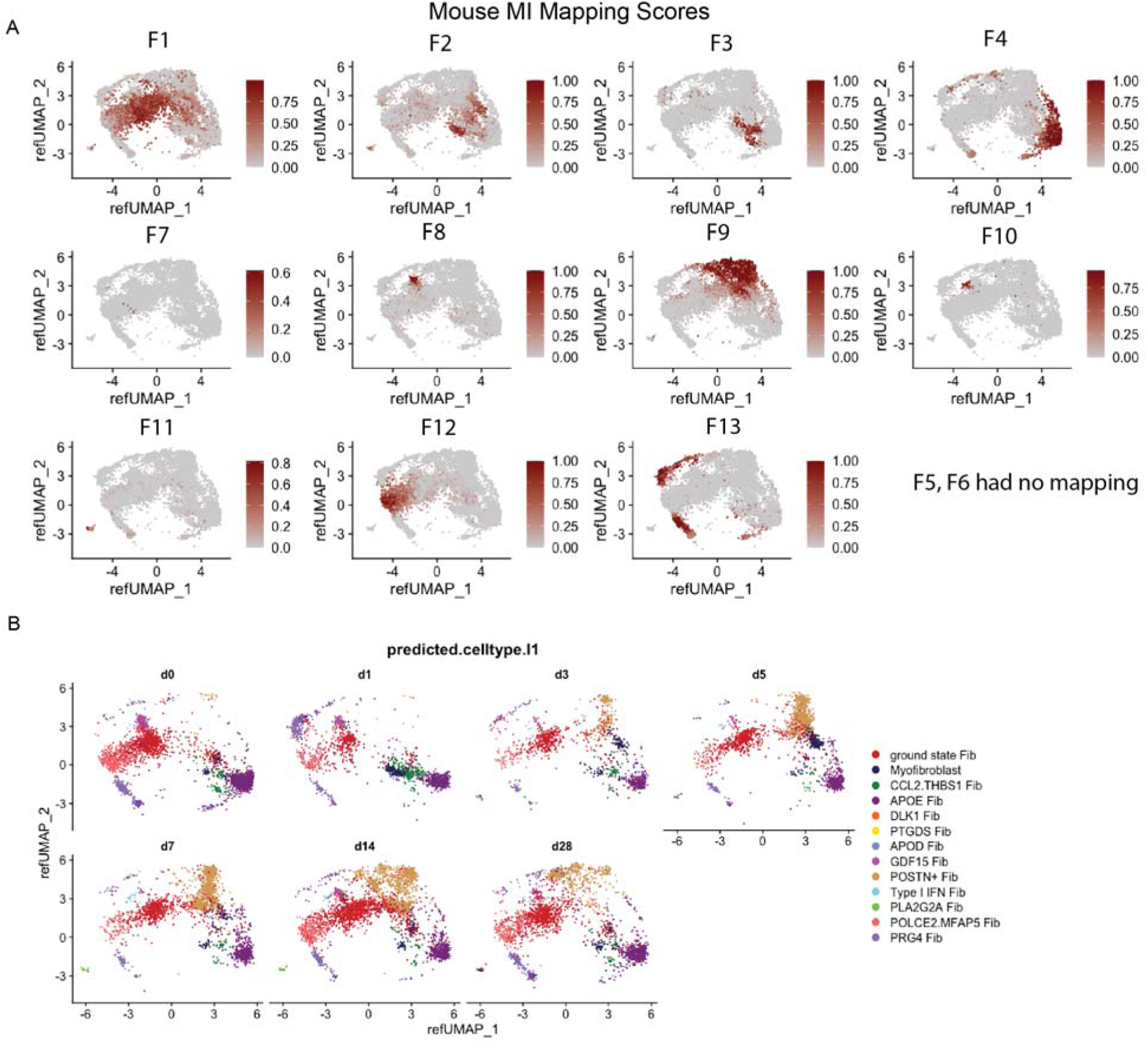
(A) Reference mapping scores for mouse MI fibroblasts onto human heart CITE-seq fibroblast UMAP embedding. (B) Reference mapped data split by MI time point in human space.

**Supplementary Figure 8.**
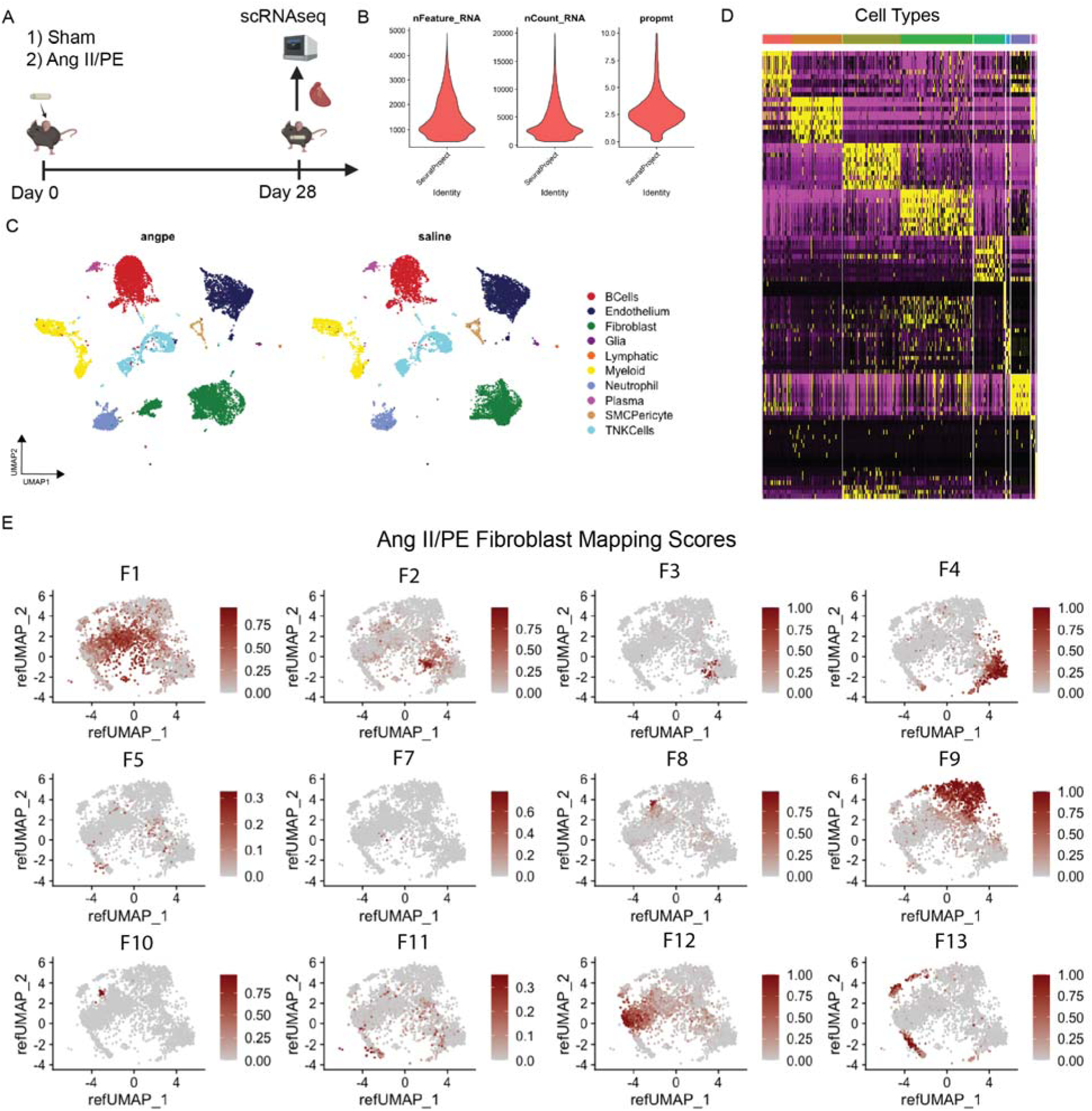
(A) Experimental design for Ang II/PE 28-day pumps for sequencing. (B) QC metrics post filtering. (C) Integrated global UMAP split by sham and Ang II/PE at day 28. (D) Heatmap of top marker genes for clusters from (C). (E) Reference mapping scores for mouse Ang II/PE and sham d28 fibroblasts onto human heart CITE-seq fibroblast UMAP embedding.

**Supplementary Figure 9.**
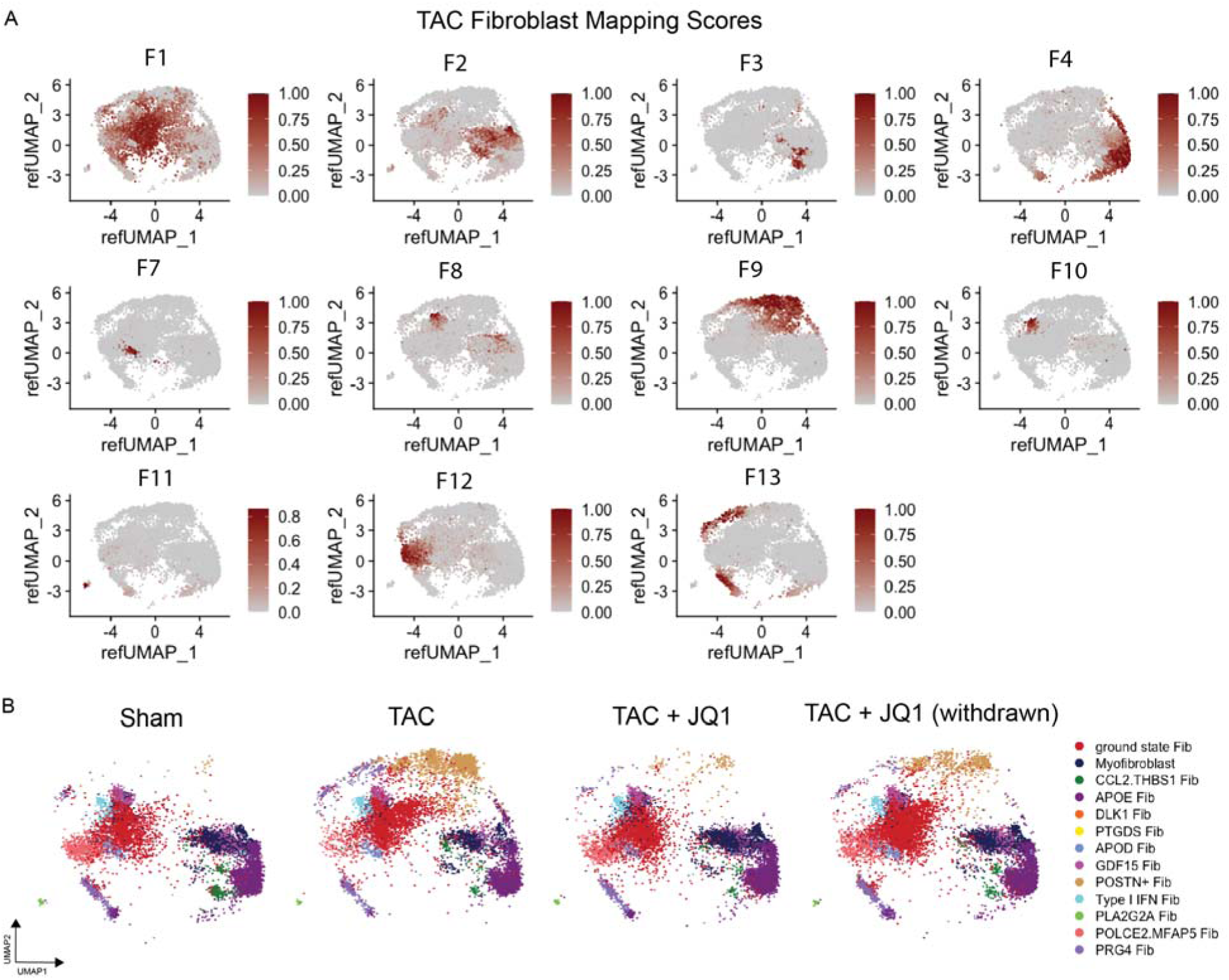
(A) Reference mapping scores for mouse TAC fibroblasts onto human heart CITE-seq fibroblast UMAP embedding. (B) Reference mapped data split by sham, TAC, TAC + JQ1 treatment, and TAC + JQ1 withdrawn in human space.

**Supplementary Figure 10.**
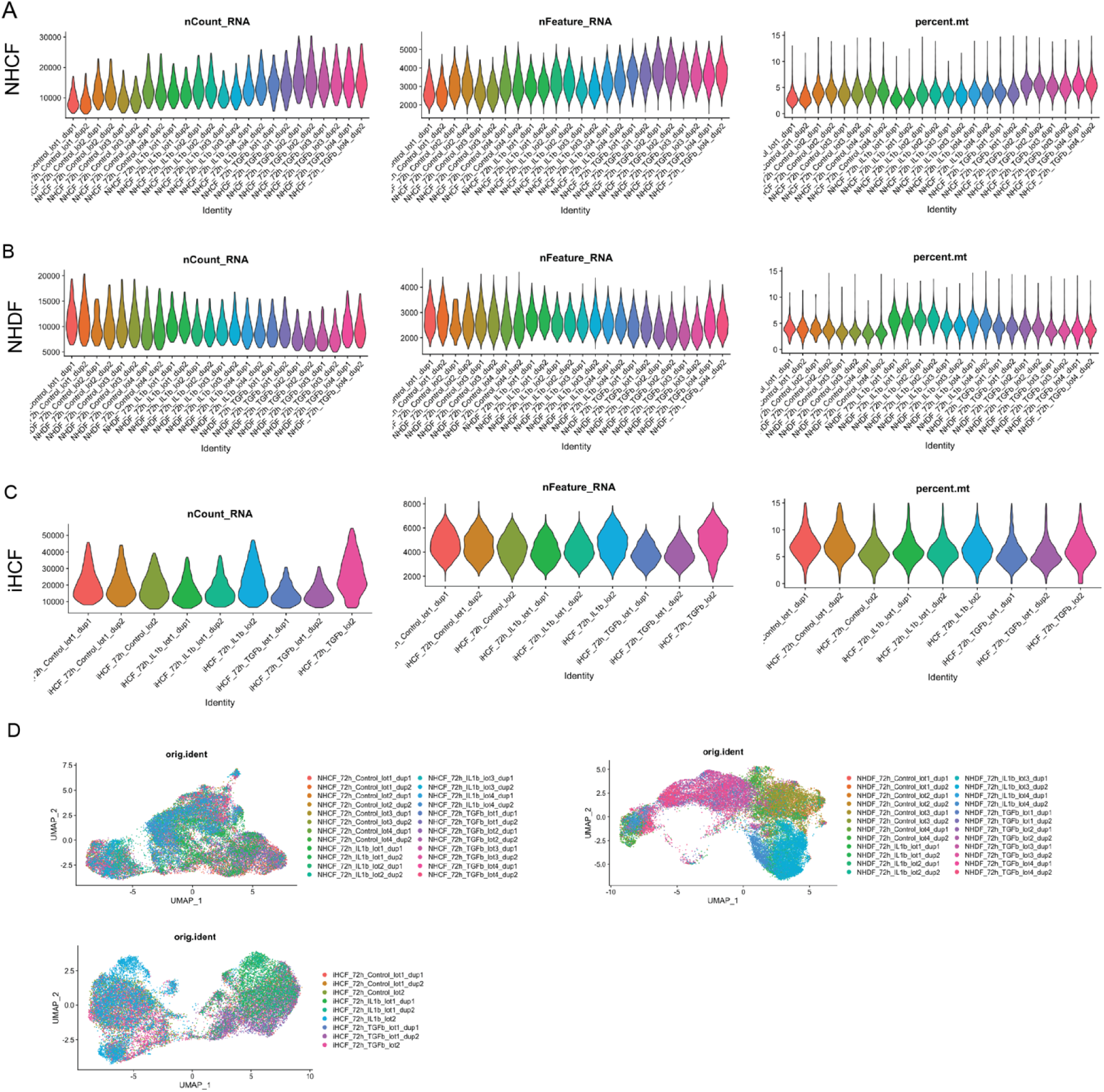
Quality control metrics post filtering grouped by each biological replicate for each experimental condition (vehicle, TGF-b, and IL-1b) in (A) NHCF, (B) NHDF, and (C) iHCF. (D) Integrated data from 3 cell lines in a UMAP embedding colored by biological replicate for each experimental condition.

**Supplementary Figure 11.**
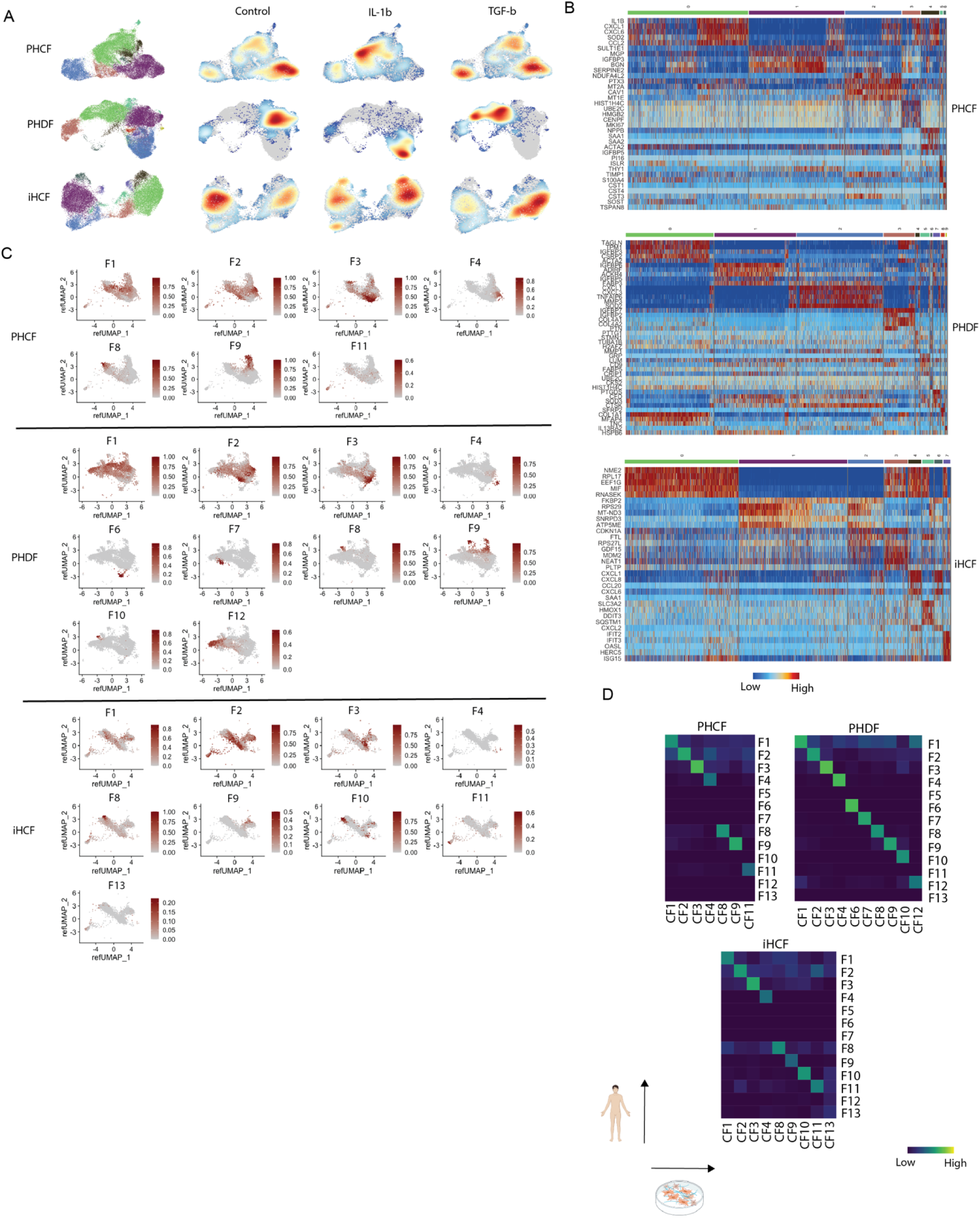
(A) UMAP embedding with separate clustering in each cell line with density plots split by experimental condition. (B) Heatmap of marker genes for each cell line for clusters in (A). (C) Reference mapping scores for *in vitro* fibroblasts onto human heart CITE-seq fibroblast UMAP embedding. (D) Heatmap of mapping scores with *in vitro* clusters on x-axis and human heart CITE-seq fibroblasts on y-axis. (E) Human, *in vivo*, and *in vitro* fibroblasts integrated in a common UMAP space colored by category. (F) Phylogeny tree clustering of categories from integrated data in (E).

**Supplementary Figure 12.**
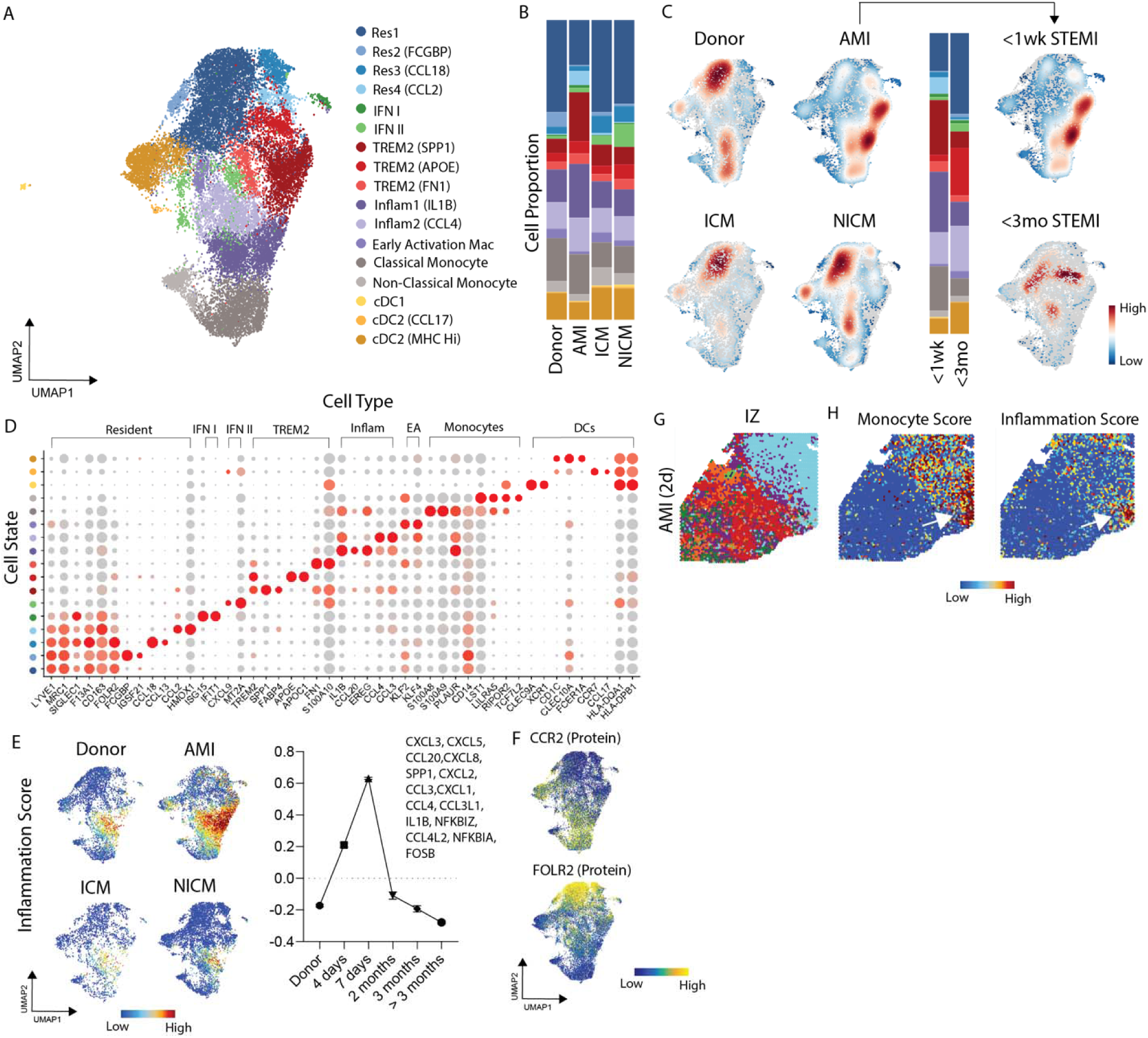
(A) UMAP embedding plot of myeloid cells with annotated cell states. (B) Myeloid cell state composition across four groups. (C) Gaussian kernel density estimation of cells across four groups (left) and split by MI time in acute MI patients with corresponding cell state composition. (D) Dot Plot of marker genes for macrophages cell states (y-axis) and grouped by cell type (x-asis). (E) Inflammation gene set score split across 4 groups and (g) grouped from time post-MI. (F) CCR2 and FOLR2 (protein) expression in UMAP embedding. (G) Spatial transcriptomic acute MI (2-day post-MI) infarct zone (IZ) sample with label transferred annotations from snRNAseq reference map. (H) Monocyte gene set score mapped into space (left) and inflammation score (right).

**Supplementary Figure 13.**
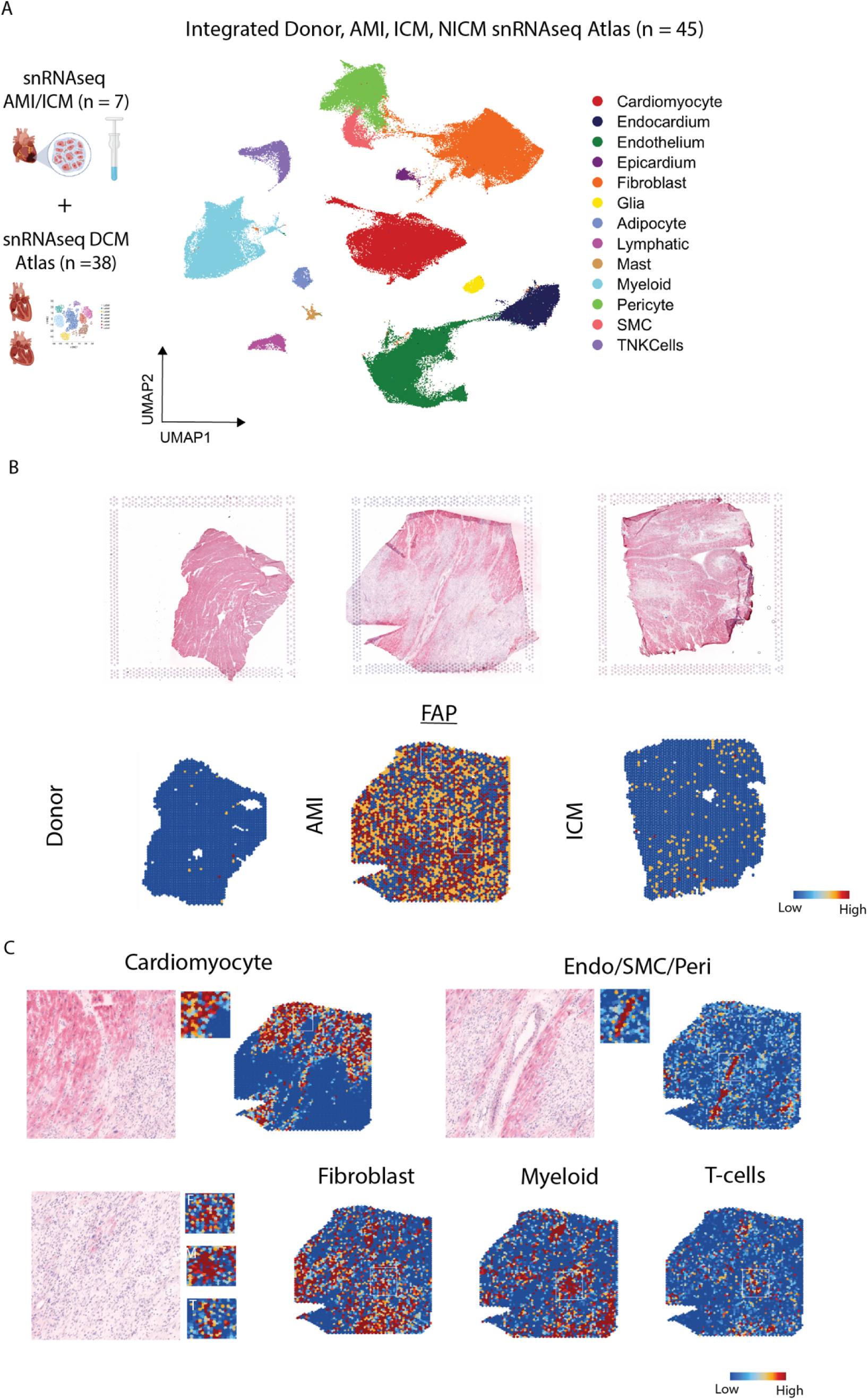
(A) UMAP embedding of integrated single nuclei RNA-seq data from donor, acute MI, ICM, and NICM. (B) HE spatial transcriptomics sections and corresponding FAP RNA expression in space. (C) Gene set signatures for cardiomyocytes, SMC/endothelium/pericytes, fibroblasts, myeloid cells, and T cells from clusters in (A) plotted in an acute MI sample.

**Supplementary Figure 14.**
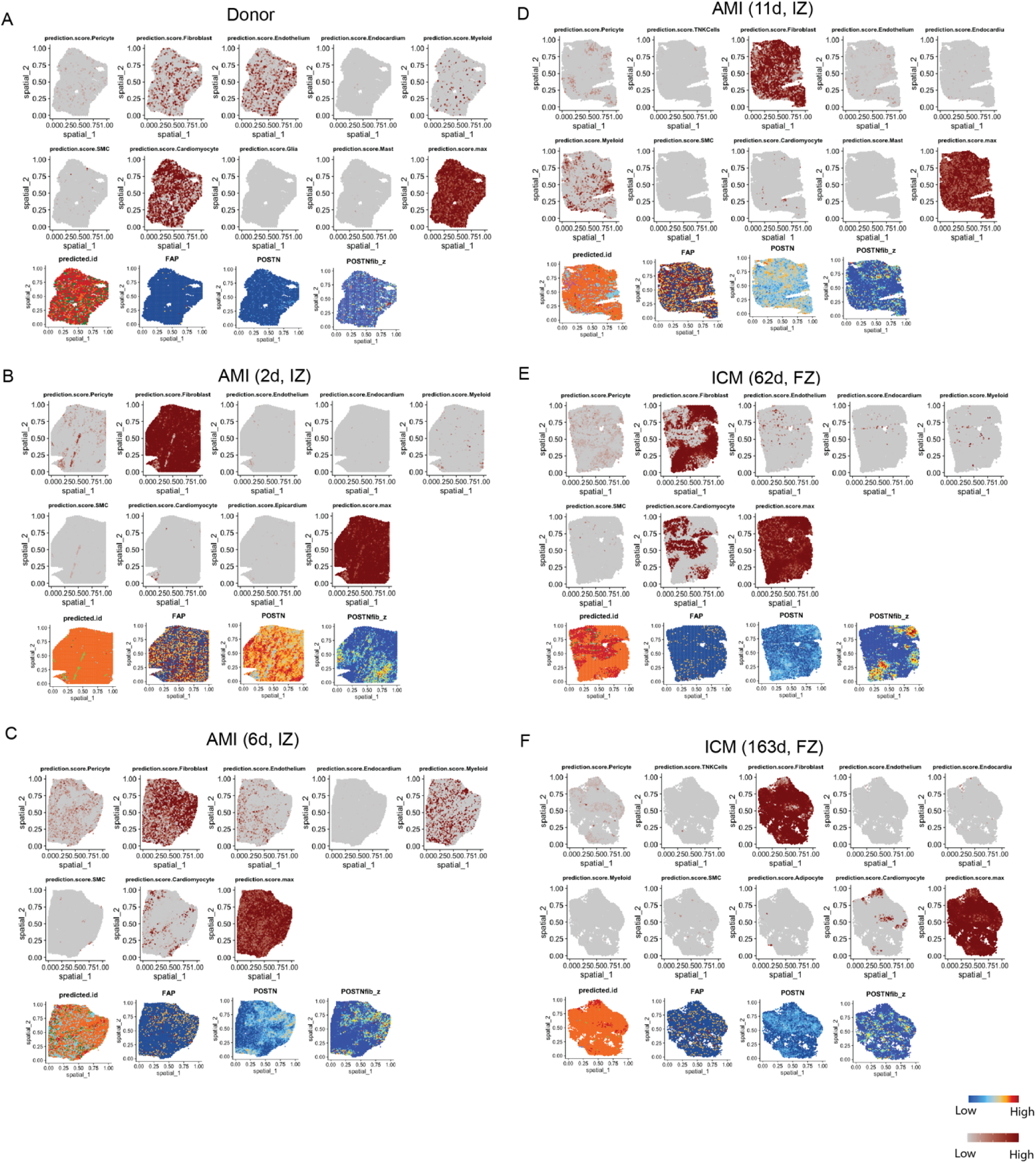
Human snRNA-seq reference map imputed prediction scores, annotations, FAP expression, POSTN expression, and F9 fibroblast gene set score in (A) donor, (B) 2d acute MI IZ, (C) 6d acute MI IZ, (D) 11d acute MI IZ, (E) 62d ICM, FZ, and (F) 163d ICM FZ sptial transcriptomic sections.

**Supplementary Figure 15.**
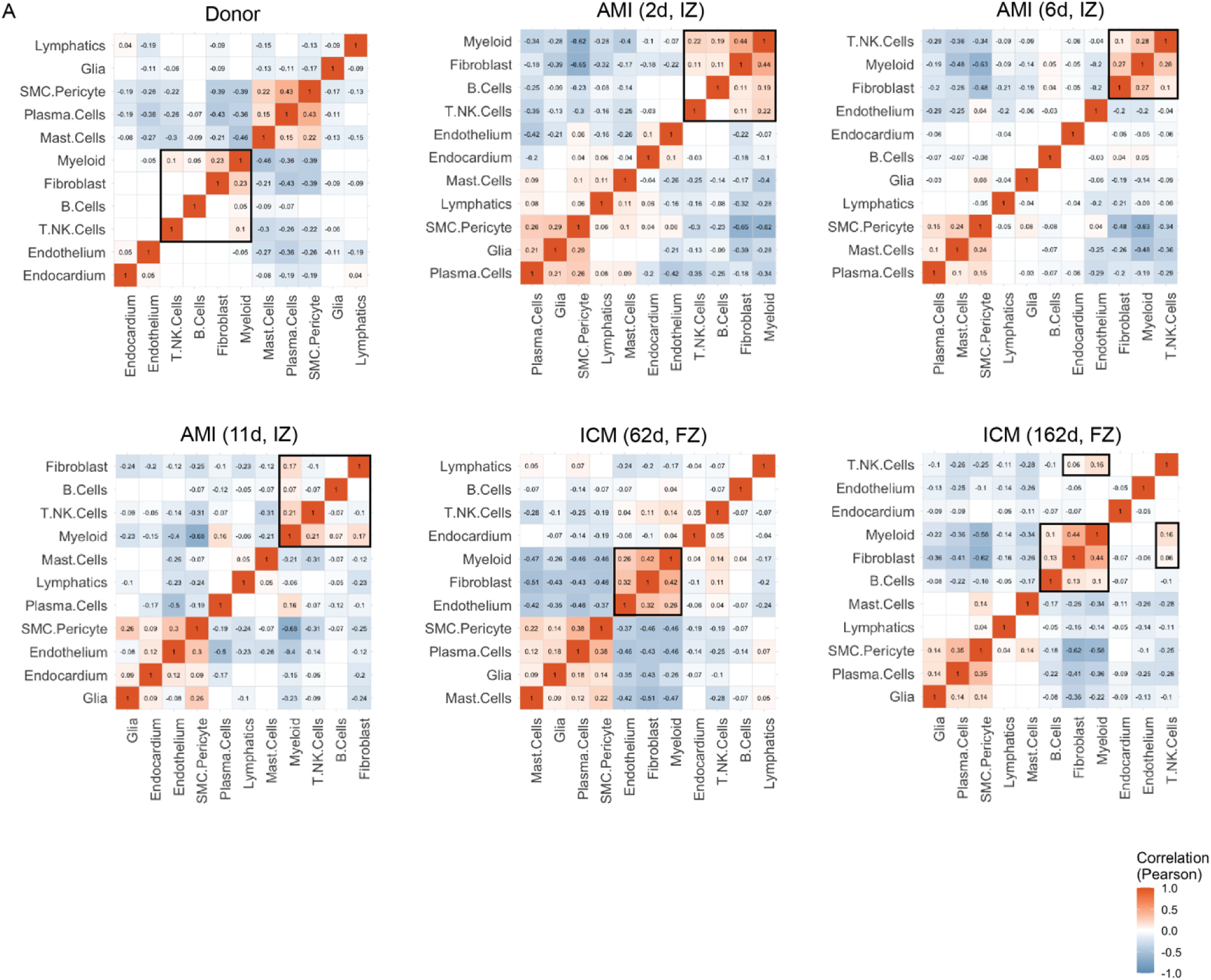
(A) SPOTlight derived cell spot deconvolution Pearson correlation coefficients for cell neighborhoods in donor, acute MI and ICM spatial transcriptomic sections.

**Supplementary Figure 16.**
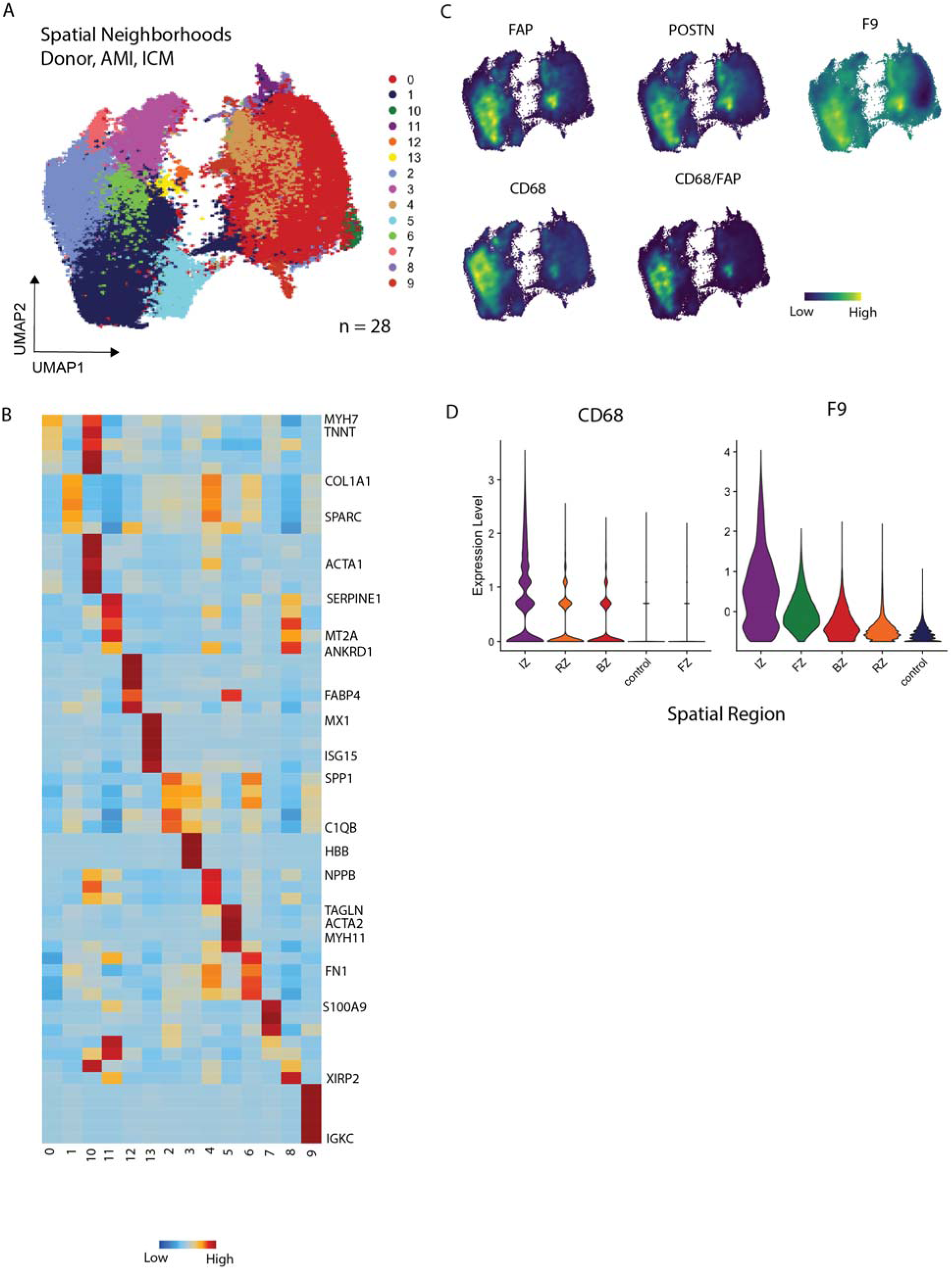
(A) Integrated UMAP embedding of n=28 spatial transcriptomic samples spatial spots. (B) Heatmap of average expression for top marker genes for spatial niches from (A). (C) FAP, POSTN, CD68 expression, CD68/FAP joint, and F9 gene set score density expression plots in integrated spatial UMAP embedding. (D) CD68 expression and F9 fibroblast gene set score violin plot split by regions.

**Supplementary Figure 17.**
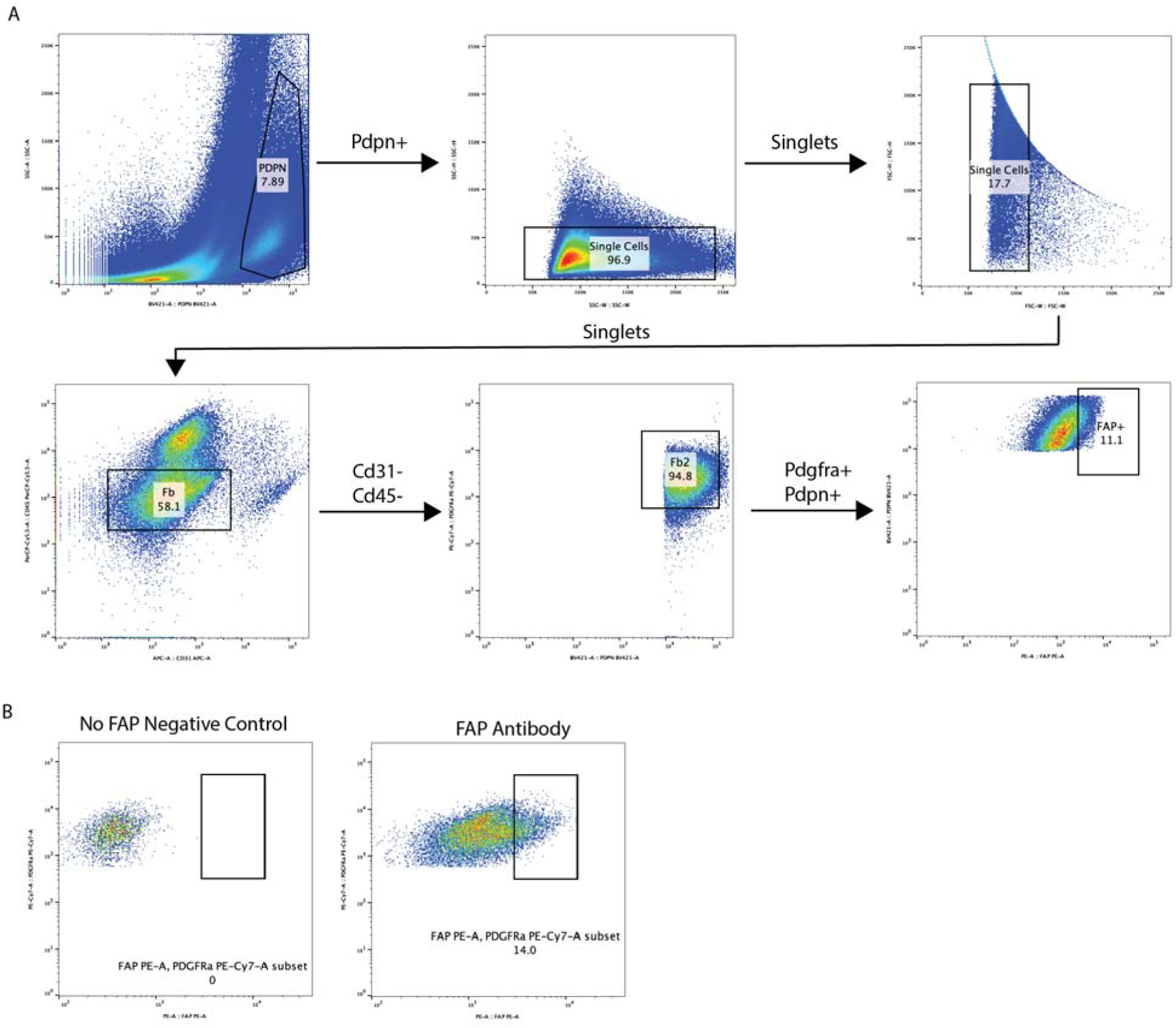
(A) Flow cytometry gating scheme for fibroblasts. (B) FAP+ gate construction with a no antibody-stained negative control.

**Supplementary Figure 18.**
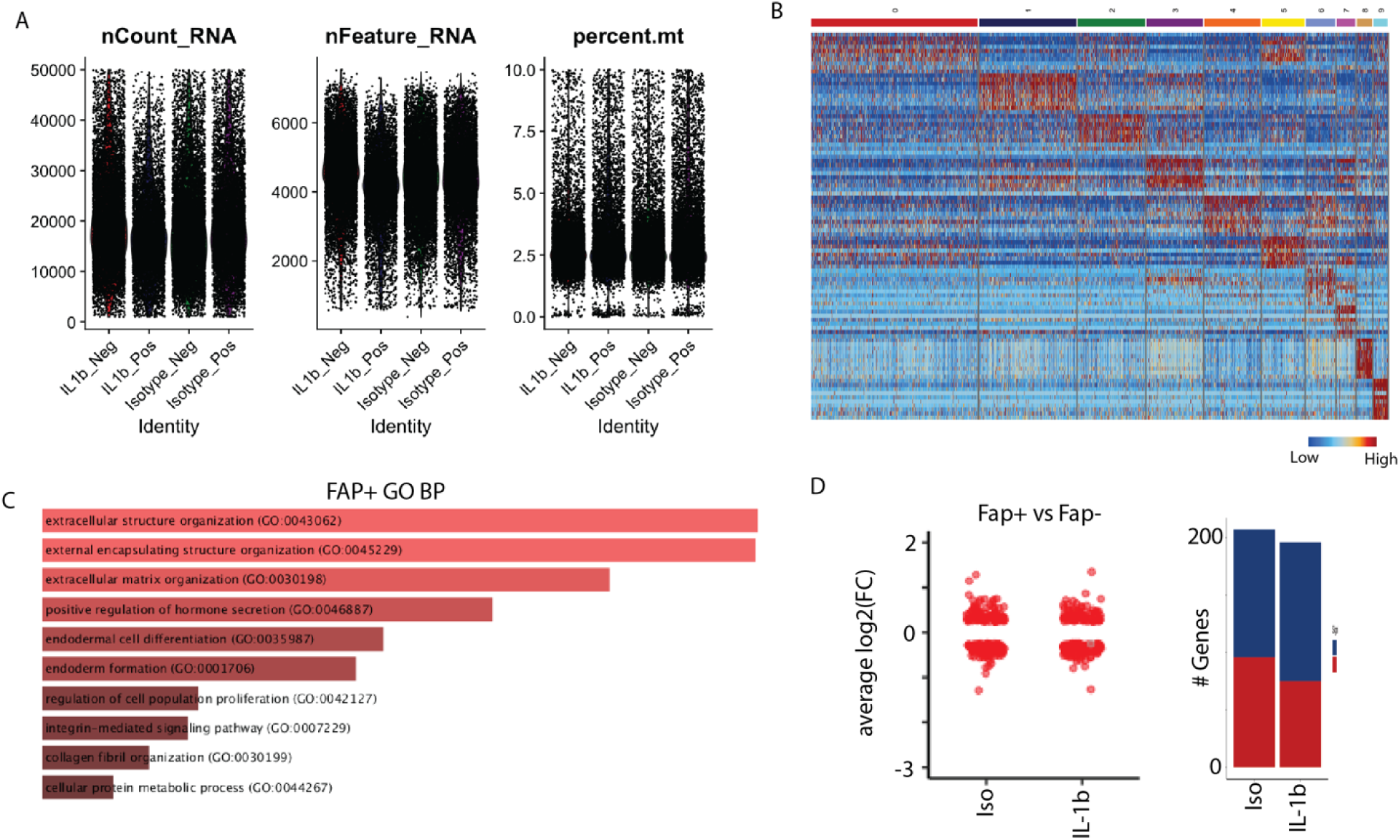
(A) QC metrics grouped by experimental condition for in vivo sorted FAP+/- fibroblasts at day 7 in Ang II/PE model with isotype/anti-IL-1b mAb treatment. (B) Heatmap of top marker genes for clusters in Fig. 7B. (C) Top GO BP pathways enriched in FAP+ fibroblasts relative to FAP- fibroblasts. (D) DE analysis between FAP+ and FAP- fibroblasts split by isotype and anti-IL-1b mAb groups dot plot (left) and number of genes up/down quantified (right).

**Supplementary Figure 19.**
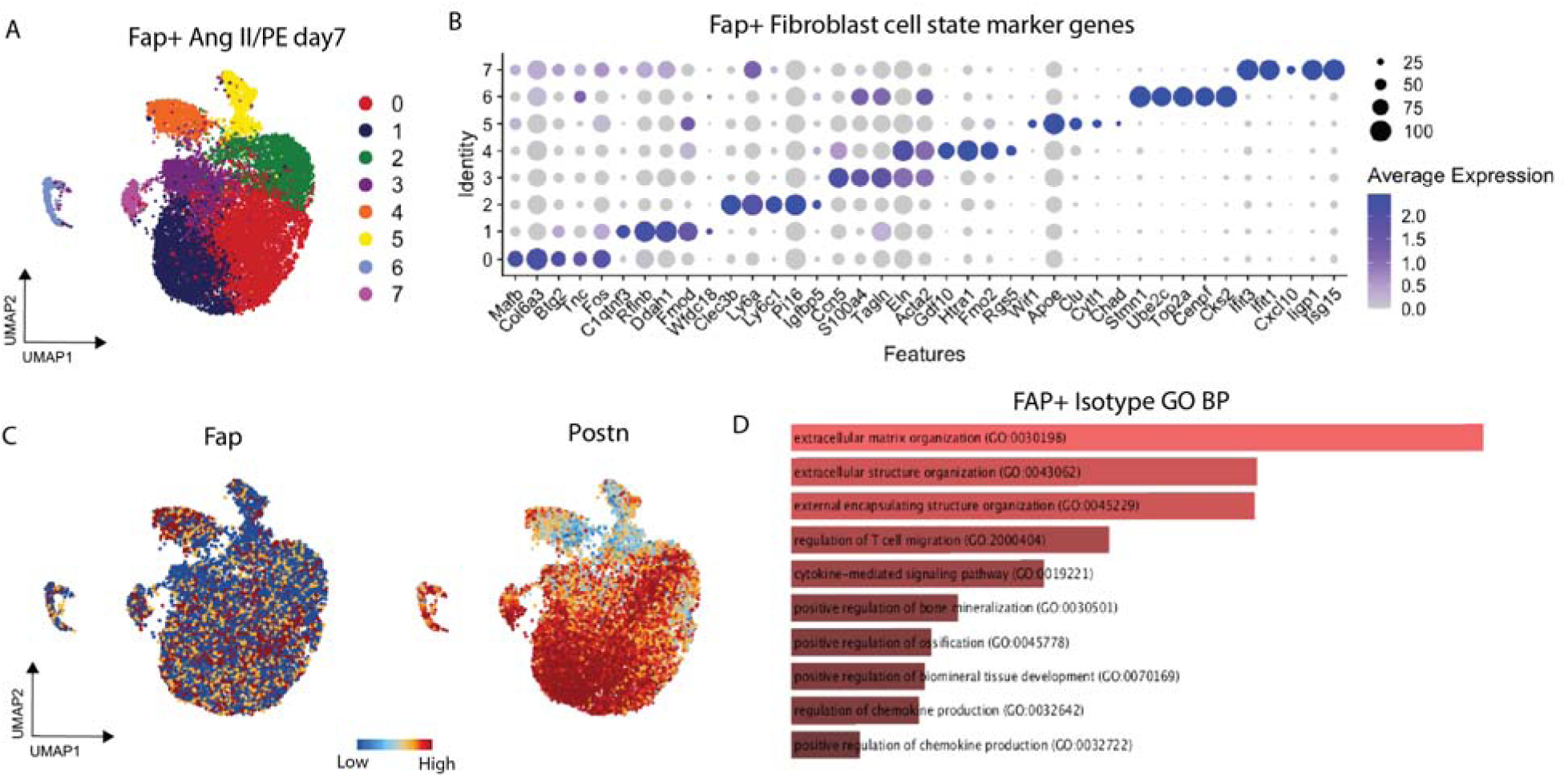
(A) Integrated UMAP with cub-clustering of FAP+ fibroblasts at day 7 in Ang II/PE. (B) DotPlot of top marker genes for FAP+ cell states from (A). (C) Fap and Postn expression in UMAP embedding. (D) GO BP pathways enriched in isotype treated FAP+ fibroblasts relative to FAP+ anti-IL-1b mAb treated mice.

